# Osteoclast-mediated resorption primes the skeleton for successful integration during axolotl limb regeneration

**DOI:** 10.1101/2022.04.27.489662

**Authors:** Camilo Riquelme-Guzmán, Stephanie L. Tsai, Karen Carreon Paz, Congtin Nguyen, David Oriola, Maritta Schuez, Jan Brugués, Joshua D. Currie, Tatiana Sandoval-Guzmán

## Abstract

Early events during axolotl limb regeneration include an immune response and the formation of a wound epithelium. These events are linked to a clearance of damaged tissue prior to blastema formation and regeneration of the missing structures. Here, we report the resorption of calcified skeletal tissue as an active, cell-driven and highly regulated event. This process, carried out by osteoclasts, is essential for a successful integration of the newly formed skeleton. Indeed, the extent of resorption is directly correlated with the integration efficiency. Moreover, we identified the wound epithelium as a major regulator of skeletal resorption, likely creating a zone of influence in which signals involved in recruitment/differentiation of osteoclasts are released. Finally, we reported a correlation between resorption and blastema formation, particularly, a coordination of resorption with cartilage condensation. In sum, our results identify resorption as a major event upon amputation, playing a critical role in the overall process of skeletal regeneration.

## INTRODUCTION

Urodele amphibians, such as the axolotl (*Ambystoma mexicanum*) are widely considered prodigies among regenerative vertebrates. The ability to regenerate different body structures, especially the limb, has driven years of scientific research aiming to understand the mechanisms underlying regeneration. The axolotl limb is a complex structure, and its regeneration requires an intricate choreography of all the cellular components. Beyond making new cells of the right type at the right place, a successful regeneration requires a functional integration of those new cells with the pre-existing tissue, a process that has not been widely studied. In particular, remains unknown how early processes impact tissue integration.

In general, regeneration progression is marked by different overlapping phases, which lead to the re-establishment of the missing limb (Sandoval-Guzmán and Currie, 2018). Two of the most critical events are the formation of the wound epithelium (WE) and the blastema (Aztekin, 2021; Tanaka, 2016). The WE is formed by migrating keratinocytes which close the wound in just a few hours (Hay and Fischman, 1961; Repesh and Oberpriller, 1978). Importantly, the WE is characterized by the absence of a basal lamina, which enhances the diffusion of important factors for regeneration (Neufeld and Aulthouse, 1986; Repesh and Oberpriller, 1978). Indeed, the WE is a major regulator of the immune response, tissue histolysis (Tsai et al., 2020), and blastema proliferation and patterning (Boilly and Albert, 1990; Ghosh et al., 2008; Han et al., 2001). Notably, several works have demonstrated that the WE is required for blastema formation and thus, regeneration (Mescher, 1976; Tassava and Garling, 1979; Thornton, 1957; Tsai et al., 2020).

The blastema is a heterogenous pool of progenitor cells arising from the various tissues at the amputation plane (Kragl et al., 2009). Among the various limb components, the connective tissue (CT) is a critical cell source for the blastema, supplying well over 40% of the cells within (Currie et al., 2016; Dunis and Namenwirth, 1977; Gerber et al., 2018; Muneoka et al., 1986). Limb CT is a conglomerate of different cell types which are found in tendons, skeleton, dermis and surrounding the skeleton (i.e. periskeleton), muscle and blood vessels. A particular case is the skeleton, where cells embedded in the skeletal matrices do not actively participate in regeneration (Currie et al., 2016; McCusker et al., 2016), instead, dermal and periskeletal cells rebuild the new skeleton (Currie et al., 2016; Dunis and Namenwirth, 1977; McCusker et al., 2016; Muneoka et al., 1986). Although the skeleton represents more than 50% of the exposed surface upon amputation (Hutchison et al., 2007), it is unclear the role the embedded cells play in the remodeling and integration of new tissue.

Undoubtedly, the skeletal system is essential for the limb, serving as a physical scaffold and allowing locomotion. In mammals, the appendicular skeleton develops by endochondral ossification, a process where a cartilage anlage is replaced by bone (Kozhemyakina et al., 2015). In axolotls, we showed that the limb skeleton is progressively ossified with growth and age (Riquelme-Guzmán et al., 2021), but retains a cartilage anlage even in the oldest specimen analyzed. Juvenile axolotls present a cartilaginous skeleton composed of chondrocytes and perichondral cells. Around the time animals reach sexual maturity, the cartilaginous skeleton is partly replaced by bone cells, adipocytes, and blood vessels during ossification. Key players in this process are osteoclasts, a myeloid-derived population, which mediates the degradation of the cartilage matrix prior to bone formation.

Osteoclasts are giant multinucleated cells with a specialized morphology adapted for skeletal resorption (Cappariello et al., 2014). Besides their role in homeostasis, osteoclasts are recruited upon bone injuries or trauma. The most studied case is fracture healing (Einhorn and Gerstenfeld, 2015); however, in the context of regeneration, osteoclasts are recruited after fin amputation in zebrafish (Blum and Begemann, 2015), and digit tip amputation in mouse (Fernando et al., 2011). In urodeles, evidence of osteoclast-mediated resorption is scarce (Fischman and Hay, 1962; Tank et al., 1976). Nevertheless, the presence of myeloid cells triggered by the amputation has been reported (Debuque et al., 2021; Leigh et al., 2018; Rodgers et al., 2020), and the participation of macrophages was shown to be critical. When macrophages were ablated, a complete halt in regeneration was reported (Godwin et al., 2013). Similar results were observed upon mouse digit tip amputation, and a specific osteoclast inhibition resulted in delayed bone resorption, wound closure and blastema formation; however, regeneration proceeded (Simkin et al., 2017).

Immune cells play an important role in histolysis, which involves the degradation of the extracellular matrix (ECM) in the vicinity of the amputation plane (Stocum, 2017), helping the mobilization of progenitor cells (Thornton, 1938a, 1938b). Histolysis is characterized by the release of proteolytic enzymes, essential for an efficient regeneration (Huang et al., 2021; Vinarsky et al., 2005; Yang et al., 1999; Yang and Byant, 1994). Histolysis is additionally controlled by the WE (Vinarsky et al., 2005), as shown by a major down-regulation of degrading enzymes upon the inhibition of WE formation (Tsai et al., 2020). Similarly, macrophage ablation resulted in a down-regulation of matrix metalloproteinases (MMPs) (Godwin et al., 2013).

Successful limb regeneration is achieved by a complete amalgamation of the regenerated structures with the mature tissues, or tissue integration. Although the regenerated limb is often considered a perfect replica of the pre-existing limb, in the last decade the fidelity of limb regeneration has been addressed by a couple of works. For instance, abnormalities due to conspecific bites were observed in 80% of larvae and 50% of adults (Thompson et al., 2014), or anomalies in over 50% of the amputated animals, such as fractures at the level of amputation or constrictions of the skeletal elements (Bothe et al., 2020). However, it is still unknown why such phenotypes are observed, and what entails successful versus unsuccessful regeneration. In this regard, regeneration-specific signals in the stump tissue could prime the limb and promote a successful integration of the newly formed structures. Indeed, in the newt *Cynops pyrrhogaster*, structural changes in the ECM of the distal humerus can be observed after an elbow joint amputation, demonstrating a correlation between ECM remodeling and proper joint regeneration as well as integration to the mature tissue (Tsutsumi et al., 2015).

With all the aforementioned evidence, we sought to assess the significance of skeletal histolysis for regeneration. We observed a rapid skeletal resorption which is carried out by osteoclasts, and we provide evidence that this process is essential for tissue integration. Moreover, we propose a role for the WE in resorption induction and a spatiotemporal coordination between resorption and blastema formation. Overall, our work provides an in-depth assessment of how a remodeling process influences the final outcome of regeneration using the axolotl limb.

## RESULTS

### Skeletal elements are resorbed upon amputation

To determine the changes in the skeleton upon amputation, we used the stable calcium-binding dyes calcein and alizarin red. These dyes label mineralized cartilage in juvenile axolotls, allowing *in vivo* imaging (Riquelme-Guzmán et al., 2021). Using 4-6 cm ST (snout-to-tail) axolotls, we amputated the zeugopod at the distal end of the calcified tissue and imaged at different days post amputation (dpa) (Fig. 1A). We observed a consistent reduction in the calcein^+^ tissue from 7 until 12 dpa. We quantified the length of the calcified tissue in both zeugopodial elements and compared them to the initial length at day 0 (Fig. 1B). Resorption initiated after 7 dpa and by 12 dpa, over 40% of the calcified radius and 60% of the calcified ulna were resorbed (length resorbed radius: 342.83 ± 95.75 µm; ulna: 770.67 ± 94.34 µm). We pooled five independent experiments and noticed an important variability between assays (Fig. 1C, each color represents an assay). The median for radius resorption is 40% and for ulna 60%; however, in several cases the calcified tissue was completely resorbed in both elements. Although an inter-assay variability was observed, intra-assay animals presented a consistent resorption ratio.

**Figure 1:**
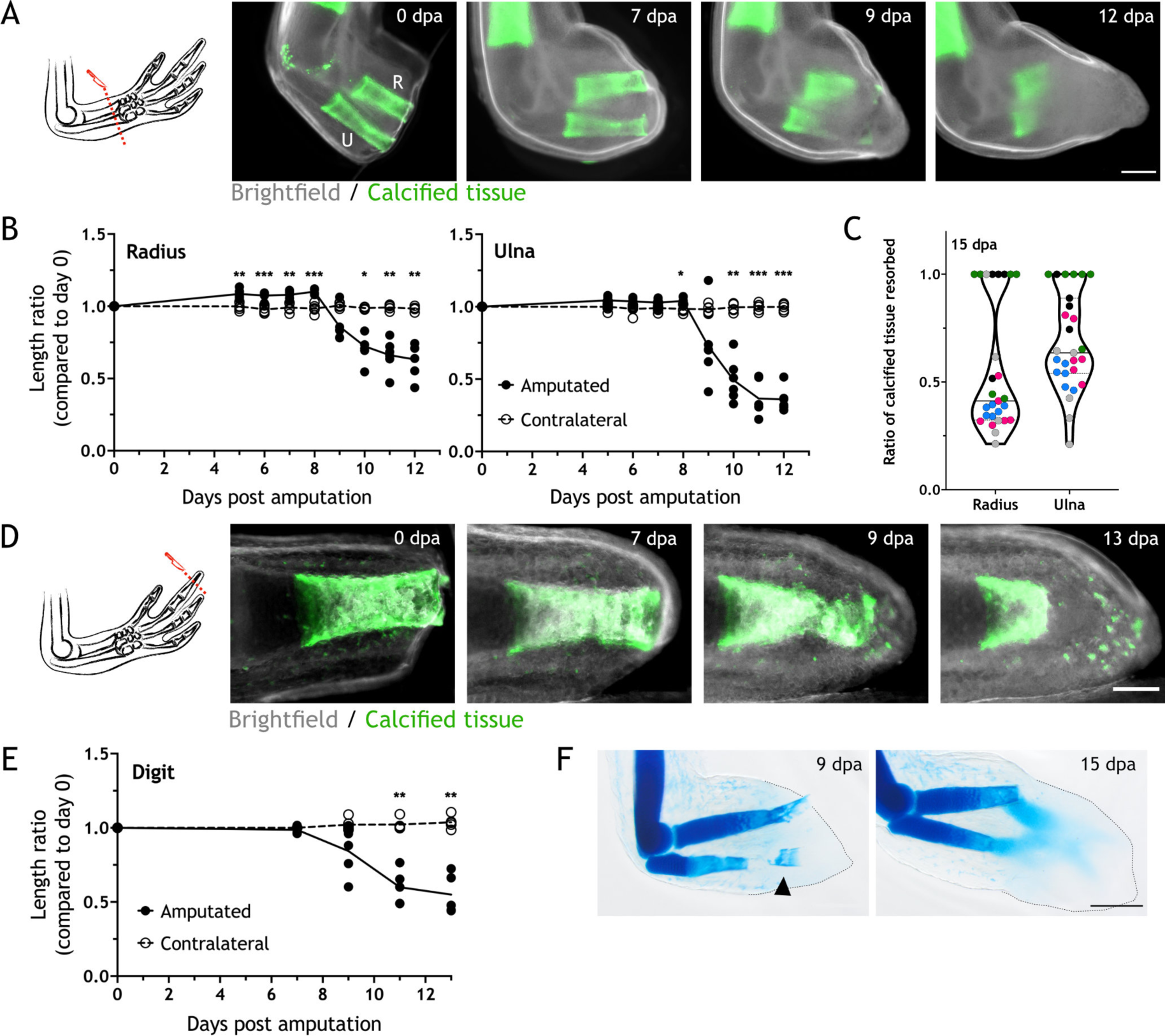
Skeletal elements are resorbed upon amputation. (A) Time course of resorption during zeugopod regeneration. Calcein-stained axolotls were amputated at the distal end of the calcified tissue (n = 6, five independent experiments). R = radius. U = ulna Scale bar: 500 µm. (B) Quantification of resorption rate in radius and ulna. Length ratio was calculated using the length at 0 dpa as a reference. Each dot represents an animal (n = 6. *** p < 0.001, ** p < 0.01, * p < 0.05, Bonferroni’s multiple comparisons test, amputated versus contralateral). (C) Quantification of resorption percentage in calcified radius and ulna among animals in different assays. Each assay is represented by a color (n = 27, five independent experiments). (D) Time course of resorption during digit regeneration. Calcein-stained axolotls were amputated at the distal end of the calcified tissue (n = 5). Scale bar: 200 µm. (E) Quantification of calcified digit resorption. Length ratio was calculated using the length at 0 dpa as a reference. Each dot represents an animal (n = 5. ** p < 0.01, Bonferroni’s multiple comparisons test, amputated versus contralateral). (F) Alcian blue staining of limbs at different dpa (n = 2). Arrowhead: broken piece of ulna. Dashed line: outline of distal limb. Scale bar: 500 µm.

Digits are a simplified platform to perform *in vivo* imaging, therefore we assessed resorption by amputating the distal end of the calcein^+^ tissue in the distal phalanx of the second digit (Fig. 1D). Similar to the zeugopod, we quantified the calcein^+^ tissue length at different dpa and revealed a similar trend in the resorptive dynamics: resorption starting after 7 dpa and receding by 13 dpa (Fig. 1E), vanishing over 50% of the calcified tissue length (320.43 ± 113.56 µm). In sum, we report resorption to be a process that occurs upon amputation of different calcified skeletal elements in the axolotl limb.

Finally, we collected limbs at 9 and 15 dpa and stained them with alcian blue (Fig. 1F). At 9 dpa, we observed resorption in both radius and ulna. Remarkably, we occasionally observed a break in the ulna (Fig. 1F arrowhead) that sometimes led to the extrusion of the skeletal fragment through the epidermis. This skeletal shedding was observed both in digit and limb amputations. At 15 dpa, resorption was finished and the condensation of the new skeleton could be observed.

### Osteoclasts are identified during skeletal resorption

Osteoclasts are specialized multinucleated cells responsible for skeletal resorption (Charles and Aliprantis, 2014). Despite their critical role in skeletal biology, osteoclasts in salamander regeneration have only been reported on the basis of morphology during salamander regeneration (Fischman and Hay, 1962; Nguyen et al., 2017; Tank et al., 1976). Therefore, we sought to identify osteoclasts during resorption using various molecular markers.

Several enzymes, such as cathepsin K (CTSK) and the tartrate-resistant acid phosphatase (TRAP) (Cappariello et al., 2014), are released by osteoclasts and are used as identifying markers. Using sections from zeugopodial amputations, we performed immunofluorescence using an anti-CTSK antibody (Fig. 2A) and TRAP enzymatic staining (Fig. 2B). CTSK^+^ cells were identified in sections at 8 dpa adjacent or inside the calcein^+^ skeleton. Similarly, TRAP^+^ cells were identified at 9 dpa. Next, to correlate osteoclast recruitment with resorption timing, we performed RT-qPCR at different dpa using specific primers for *Trap*, *Ctsk* and *Dcstamp* (dendritic cell-specific transmembrane protein, involved in osteoclast multinucleation). The RNA relative content for the three markers behaved similarly: a sharp increase was observed, reaching a peak at 9 dpa before rapidly decrease to almost basal levels at 15 dpa (Fig. 2C).

**Figure 2:**
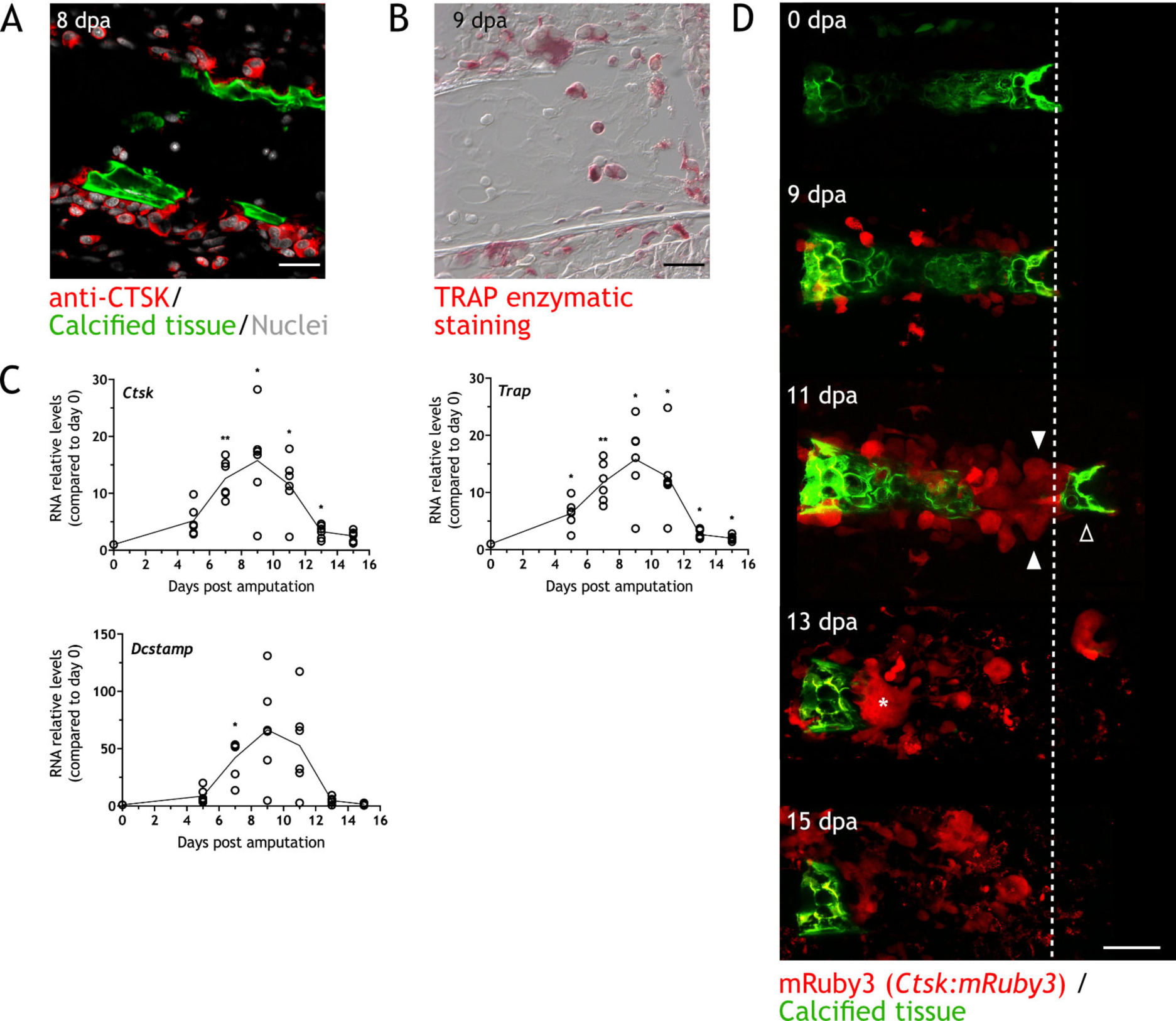
Osteoclasts are identified during skeletal resorption. (A) Apotome image of IF for anti-CTSK (red) in zeugopod section at 8 dpa (n = 2). Calcein was used for calcified cartilage labelling (green) and Hoechst for nuclear staining (white). Scale bar: 50 µm. (B) TRAP enzymatic staining in zeugopod section at 9 dpa (n = 2). Scale bar: 50 µm. (C) RT-qPCR for T*rap*, C*tsk* and D*cstamp* at different dpa upon zeugopodial amputation. Solid line represents mean, each dot is an animal (n = 6. ** p < 0.01, * p < 0.05, Bonferroni’s multiple comparisons test, each time point versus 0 dpa). (D) *In vivo* confocal imaging of *Ctsk:mRuby3* (red) upon digit amputation (n = 3, two independent experiments). Calcein was used for calcified cartilage labelling (green). Image represent a maximum intensity projection of 10 images (3 µm interval). White arrowhead: mRuby3^+^ cells (osteoclasts). Black arrowhead: break in the skeletal tissue. Dashed line: amputation plane. Asterisk: Multinucleated osteoclast. Scale bar: 100 µm.

To assess osteoclast spatiotemporal dynamics *in vivo*, we developed a *Ctsk:mRuby3* and *Ctsk:eGFP* transgenic lines, which express the fluorescent protein *mRuby3* or *eGFP* under the control of *Ctsk* promoter from zebrafish. Using *Ctsk:mRuby3* animals, we followed resorption in digits with confocal microscopy (Fig. 2D). At 0 dpa, the tissue was devoid of mRuby3^+^ cells. At 9 dpa, mononuclear-like cells were observed in the periphery of the calcified phalanx. These cells increased in numbers and size at 11 dpa (white arrowheads). A break in the phalanx (black arrowhead) was seen at this timepoint. At 13 dpa, most of the phalanx was resorbed and mRuby3^+^ cells were scattered throughout the sample. A giant multinucleated cell was observed next to the calcified tissue (asterisk Fig. 2D). Finally, between 13 and 15 dpa, resorption was completed and mRuby3^+^ cells vacated the space, some showing signs of apoptotic puncta. Although most osteoclasts were multinucleated, we also observed mononuclear cells. Whether this indicates differences in osteoclast biology between axolotls and other model organisms, is not yet determined. Nevertheless, by utilizing different approaches, we demonstrated the presence and participation of osteoclasts in the regeneration-induced resorption.

### Zoledronic acid treatment inhibits osteoclast-mediated skeletal resorption

To assess the effect of osteoclast inhibition on regeneration, we treated animals with the osteoclast inhibitor zoledronic acid (zol). Zol is a potent bisphosphonate, used in the treatment of osteoporosis. It is internalized by osteoclasts, preventing protein prenylation and consequently their intracellular localization and function (Dhillon, 2016), which could lead to apoptosis (Clézardin, 2013). By serial intraperitoneal injections of 200 µg/kg of zol every 3 days, we evaluated the effect of osteoclast inhibition by imaging the length of the skeletal elements at different dpa. Zol treatment inhibited resorption as seen at 12 dpa (Fig. 3A), since most of the calcified tissue remained intact when compared to the untreated control and vehicle. Quantification of both radius and ulna lengths at different dpa revealed a significant difference between the radius or ulna in zol-treated animals compared to the controls at 11, 12 and 15 dpa (Fig. 3B).

**Figure 3:**
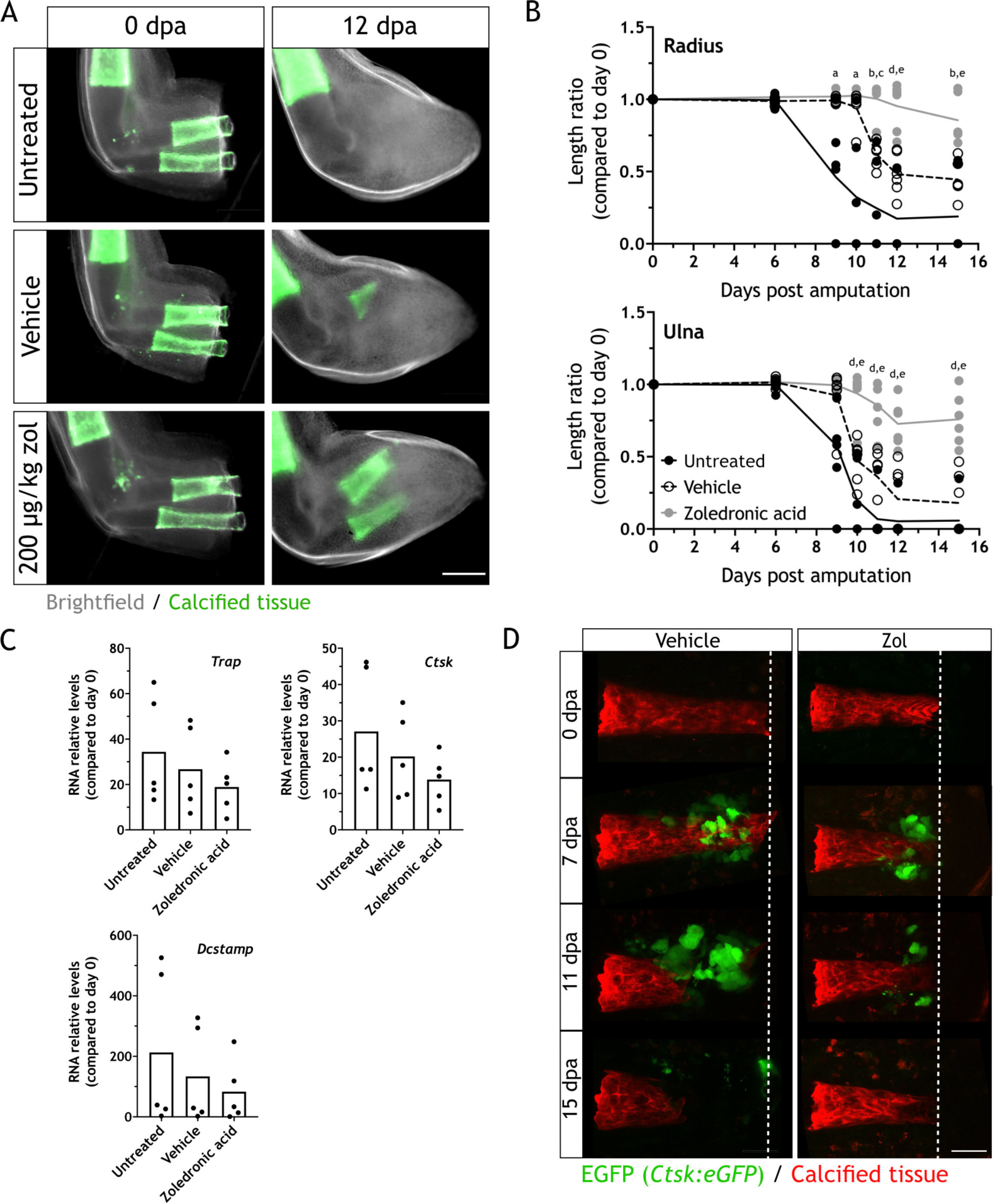
Zoledronic acid treatment inhibits osteoclast-mediated skeletal resorption. (A) Time course of resorption during zeugopod regeneration upon zoledronic acid treatment (zol) (n = 6, three independent experiments). Calcein-stained axolotls were amputated at the distal end of the calcified tissue. Scale bar: 500 µm. (B) Quantification of resorption rate in radius and ulna upon zol treatment. Length ratio was calculated using the length at 0 dpa as a reference. Each dot represents an animal (n = 6. a: p < 0.05 uninjected vs. zol, b: p < 0.01 uninjected vs. zol, c: p < 0.001 vehicle vs. zol, d: p < 0.001 uninjected vs. zol, e: p < 0.01 vehicle vs. zol, Tukey’s multiple comparisons test). (C) RT-qPCR for *Trap*, C*tsk* and D*cstamp* at 9 dpa upon zol treatment. Each dot represents an animal (n = 5, Tukey’s multiple comparisons test). (D) *In vivo* confocal imaging of *Ctsk:eGFP* (green) upon digit amputation (n = 4, two independent experiments). Alizarin red was used for calcified cartilage labelling (red). Image represent a maximum intensity projection of 15 images (3 µm interval). Scale bar: 50 µm.

Furthermore, we measured the relative RNA content of *Ctsk*, *Trap* and *Dcstamp* at 9 dpa in each condition. No significant difference was observed for the three markers (Fig. 3C), although the mean for zol-treated samples was smaller in each case. Our results suggested that zol treatment mainly results in a consistent inhibition of osteoclast function. Consequently, we performed *in vivo* imaging of digit regeneration upon zol treatment in the *Ctsk:eGFP* transgenic line. When resorption was inhibited by zol treatment, we observed a reduction in the number of eGFP^+^ cells (Fig. 3D). Although present, these cells did not seem to resorb the calcified tissue. Therefore, zol treatment inhibits osteoclast-mediated resorption, but it does not result in their complete ablation.

### Skeletal resorption is necessary for a successful integration of the regenerated structure

To assess the importance of resorption, we followed the zol-treated animals until 45 dpa. At this stage, limbs are fully formed but they have not reached yet full size (Tank et al., 1976). Looking at the gross morphology, resorption inhibition did not halt regeneration, as zol-treated animals were able to form a new limb. We assessed integration by sequentially staining with calcium binding dyes of different color, we distinguished the original calcification (calcein^+^) from the calcification of regenerated skeletal elements (alizarin red^+^) (Fig. 4A, left panel). In contralateral limbs, alizarin red staining showed new calcification from the calcein staining at 0 dpa. Comparatively, we observed no calcein^+^ tissue in the untreated animals, indicating a full resorption of the calcified tissue in the radius and ulna. The alizarin red^+^ region demonstrated regeneration of the skeleton. In zol-treated animals, at least half of the calcified region was calcein^+^/alizarin red^+^, confirming resorption inhibition. Interestingly, we observed a faulty morphology in the radius from the zol-treated animal (arrowhead Fig. 4A, left panel). To gain a better insight into the morphology of the regenerated zeugopod, we collected those limbs and stained them with alcian blue (Fig. 4A, right panel). The zol-treated limb showed a clear failure in the integration of the newly formed cartilage, especially in the radius. The new tissue lacked a seamless connection to the stump part, presenting an angulated morphology (black arrowhead). In the ulna, heterotopic cartilage formation was seen (asterisk). Surprisingly, the skeletal elements of untreated regenerating animals also showed imperfect morphology, even though the calcified areas were fully resorbed. Both radius and ulna were restored as one complete unit, but with an irregular interphase between the stump and regenerated part, observed as a narrowing in the mid-diaphysis (black arrowhead, Fig. 4A).

**Figure 4:**
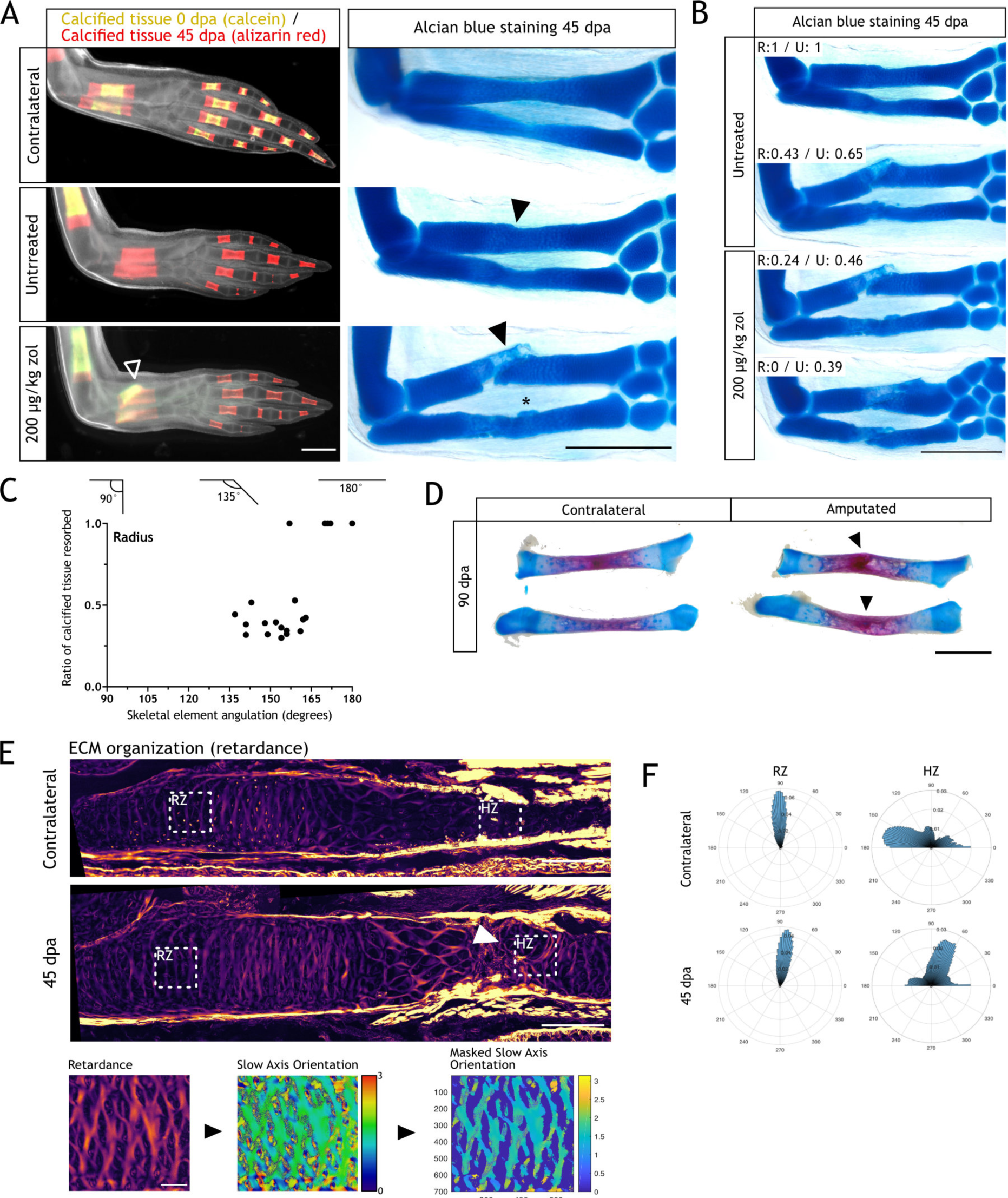
Resorption inhibition does not halt regeneration but results in an integration failure of the newly formed skeleton. (A) *In vivo* calcein / alizarin red staining (left panel) and alcian blue staining (right panel) in zol treated limbs at 45 dpa (n = 6). Arrowheads: integration failure in skeletal elements. Asterisk: heterotopic cartilage formation in ulna. Scale bar: 1 mm. (B) Alcian blue staining in zol treated limbs at 45 dpa (n = 6). Resorption rate for radius and ulna is specified for each case. Scale bar: 1 mm. (C) Quantification of angulation at the stump-regenerated interphase in untreated radii at 45 dpa. Angulation schematic is shown on top of the graph (n = 22, 4 independent experiments). (D) Alcian blue/alizarin red staining of zeugopodial elements at 90 dpa. Arrowhead: stump-regenerated interphase. Scale bar: 2 mm. (E) Upper panel: retardance image from unamputated and 40 dpa ulna. RZ (resting zone) and HZ (hypertrophic zone) squares represent the quantification areas (n = 7 for unamputated, n = 9 for amputated). Arrowhead: disorganized interphase. Scale bar: 200 µm. Lower panel: quantification flow chart. The mask was created using the retardance image to quantify only ECM components, and applied to the slow axis orientation image to determine the orientation of the ECM components at each pixel. In the masked orientational field, the cellular regions are shown in dark blue for visualization purposes but their orientational values were excluded from the analysis in F. Scale bar: 50 µm. (F) Histograms showing the orientation of the ECM components at each pixel in RZ or HZ for the unamputated or 40 dpa ulna (n = 7 for unamputated, n = 9 for amputated). Angles are shown in degrees.

Among the control and zol-treated limbs, we found different rates of resorption (Fig. 4B). A correlation between resorption rate and integration efficiency could be observed, particularly for the radius. In a zol-treated animal, with null resorption in the radius (R:0), the distal end of the stump part and the proximal end of the regenerated skeleton failed to meet. The regenerated skeleton formed at an adjacent plane, therefore lacking continuity with the pre-existing skeleton. Moreover, a second condensation zone was seen, as cartilage also formed distal to the un-resorbed tissue.

To consistently quantify the integration success, we analyzed only animals undergoing normal, undisturbed regeneration. This way, with less severe phenotypes and the skeletal elements regenerating as one unit, it was simpler to correlate resorption to integration. The angle of the regenerated skeletal elements to the stump skeletal element was measured. In 18/22 untreated animals, their radii were not fully resorbed and presented different degrees of angulation, between 135° to 165°, at the stump- regenerated interphase (Fig. 4C). These results show that even in the best-case scenario, limb regeneration in the axolotl can lead to an imperfect outcome.

We assessed whether the faulty integration was resolved at later stages. We collected limbs at 90 dpa and stained them with alcian blue/alizarin red. In 6/6 limbs screened, in which the resorption rate was over 50% for both elements, we could still observe a faulty integration of both radius and ulna (arrowheads, Fig. 4D). This imperfect integration was identified by an angulation at the stump- regenerated interphase, similar to what we reported at 45 dpa.

As resorption has a clear impact on skeletal integration during regeneration, we sought to analyze the ECM organization and its changes at the stump-regenerated interphase using quantitative polarization microscopy (LC-PolScope) (Oldenbourg, 1996). We used limb sections from normally regenerated animals for our analysis. By looking at the ECM organization (retardance image), we observed a clear difference in the hypertrophic zone (HZ) of regenerated ulnas when compared to the contralateral limb (arrowhead, Fig. 4E). We believe that this region in the HZ corresponded to the stump- regenerated interphase. Next, using the retardance image, we created a digital mask that allowed us to quantify the orientation of the ECM components using the slow axis orientation image (Fig. 4E, lower panels). We defined two regions, the HZ, where the interphase is found, and the resting zone (RZ), a control region proximal to the amputation plane. We generated a histogram representing the angle distribution in each zone. We observed that the HZ in the contralateral ulnas presented a parallel organization of the ECM respect to the proximo-distal axis, while the regenerated presented a shift in the organization, with the ECM fibers arranged perpendicularly. The RZ remained unchanged in both samples sets (Fig. 4F). This result shows that the regenerated ECM does not recapitulate the original structure, and supports the idea that skeletal regeneration is not completely efficient in the axolotl.

Altogether, these results show the importance of resorption during skeletal regeneration and requirement for integration of the regenerated tissue. However, these results also highlight that even in normal conditions, the regenerated skeleton does not recapitulate the smooth structure seen pre-amputation.

### The wound epithelium is involved in resorption induction

A previous report showed that the WE is critical for inflammation and tissue histolysis (Tsai et al., 2020). When the WE formation was prevented by mechanically closing the wound with stump tissue, in a so-called full skin flap surgery (FSF), *Ctsk* expression was absent at 5 dpa compared to control. This suggests potential defects in skeletal resorption and a role of the WE in its induction. Therefore, we sought to evaluate the role of the WE in skeletal resorption.

Given the technical difficulty of this surgical procedure, we used 14 cm ST animals, similar to the previous reported model (Tsai et al., 2020). We amputated the limbs prior to FSF surgery, and followed them until 15 dpa. Using *in vivo* imaging, we observed an inhibition in resorption in FSF samples by comparing calcified tissue length to the control limb (arrowhead Fig. 5A). Next, we collected the limbs and performed alcian blue/alizarin red staining. In 7 out 9 control samples, we observed a clear degradation in the distal end of the skeletal elements (black arrowhead Fig. 5B), while no or limited resorption was observed in the FSF limbs. To further confirm resorption inhibition, we collected limbs at 9 dpa and performed ISH for *Ctsk*. We saw a significant reduction of the *Ctsk* staining in FSF sections (Fig. 5C), suggesting the WE plays a role in the recruitment or differentiation of osteoclasts.

**Figure 5:**
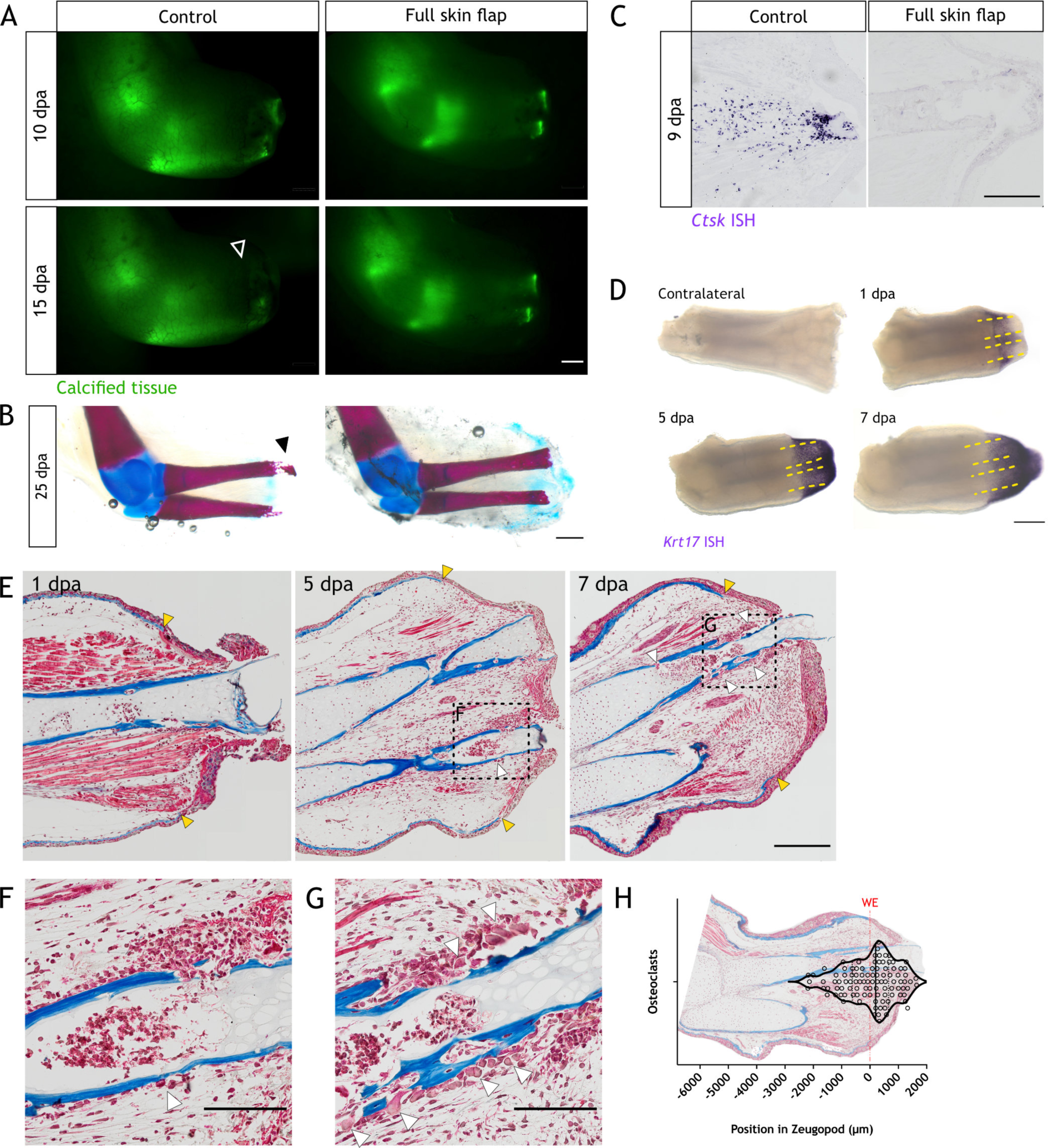
The wound epithelium is involved in resorption induction. (A) Time course of resorption during zeugopod regeneration upon full skin flap surgery (n = 9). Calcein- stained axolotls were amputated at the distal end of the calcified tissue. Arrowheads: resorption in control cases. Scale bar: 1 mm. (B) Alcian blue/alizarin red staining of limbs at 25 dpa after full skin flap surgery (n = 9). Arrowhead: resorption of distal radius. Scale bar: 1 mm. (C) ISH for *Ctsk* in limb sections at 9 dpa after full skin flap surgery (n = 3 for control, n = 4 for FSF). Scale bar: 500 µm. (D) WISH for *Krt17* in limbs upon zeugopod amputation at different dpa (n = 3). Dashed lines: skeletal elements position. Scale bar: 500 µm. (E) Masson’s trichrome staining from limb sections upon zeugopod amputation at different dpa (n = 3). Yellow arrowheads: beginning of wound epithelium. White arrowheads: osteoclasts. (F) Inset from (E) 5 dpa. Scale bar: 200 µm. (G) Inset from (E) 7dpa. White arrowheads: osteoclasts. Scale bar: 200 µm. (H) Quantification of position of osteoclasts in zeugopod at 7 dpa. Each dot represents an osteoclast. Position of WE is shown with a red line. Image of a quantified section shows the position of osteoclasts in the sample (n = 101, 3 independent experiments).

By using 14 cm ST animals, we demonstrate that resorption occurs also when skeletal elements in the limb are undergoing ossification. Limbs of older animals are opaquer and becomes harder to quantify the length of the calcified tissue, thus, we performed micro-CT scans in limbs of animals 16 cm ST (Fig. supplement 1). We confirmed a significant resorption of ossified elements in a slightly extended, but conserved time window than in small animals.

To evaluate whether the WE position might determine the region of resorption initiation, we spatially correlated resorption and the WE. We performed WISH for *Krt17,* which labels cells in the basal layers of the WE (Leigh et al., 2018). We observed a clear labelling of the WE from 1 to 7 dpa (Fig. 5D). At 1 and 5 dpa, we observed that at least 1/3 of the skeletal elements (yellow dashed lines) were covered by the WE, which could account for over 50% of the tissue that will be resorbed. Morphologically, the WE is characterized by the absence of a basal lamina (Neufeld and Aulthouse, 1986; Repesh and Oberpriller, 1978; Tsai et al., 2020), hence we used this feature to correlate the WE and resorption. We collected and sectioned limbs at 1, 5 and 7 dpa, and performed Masson’s trichrome staining. The lack of a basal lamina was observed by the absence of collagen staining in blue (yellow arrowheads, Fig. 5E). Osteoclasts could be identified by their multiple nuclei and morphology. As expected, we did not observe any osteoclast at 1dpa. At 5 dpa, we identified the first infiltration of the skeletal elements, albeit a small number of osteoclasts (white arrowheads, Fig. 5E, F). Finally, we saw pronounced infiltration at 7 dpa, including the presence of osteoclasts (white arrowheads, Fig. 5E, G). Notably, most of these cells were located in the proximity of the WE. To evaluate the location of the osteoclasts, we defined a region by drawing a line between the edges of the WE in the sections stained and we mapped the position of each osteoclast at 7dpa (Fig. 5H). Most of the osteoclasts were located in the region covered by the WE, i.e. in the distal part of the skeletal elements. In the more proximal regions of the zeugopod, we did not observe any osteoclasts.

In sum, our data suggest that the WE plays a role in both osteoclasts recruitment and/or differentiation, as well as in generating a zone of influence which determines resorption initiation in the distal regions of the skeletal elements.

### Identification of candidates involved in osteoclast recruitment and/or differentiation

To identify possible candidates involved in osteoclast recruitment and/or differentiation, we curated a published RNA-seq dataset in which FSF surgery was performed (Tsai et al., 2020). In that work, three different populations from blastemas at 5 dpa were isolated: dividing cells (4N), non-dividing cells (2N) and epithelial cells (EP). We first checked which fraction was enriched for transcripts associated with osteoclast function at 5 dpa. As expected, non-dividing cells (2N) were enriched for osteoclasts genes (*Trap, Traf6, Rank, Ocstp, Nfatc1, Dcstamp, Ctsk, Csfr1*) (Fig. 6A). Moreover, in the 2N fraction at 5 dpa, most of those transcripts were up-regulated compared to day 0, and down-regulated in FSF limbs at 5 dpa (Fig. 6B). This analysis supports our previous results, in which we observed an inhibition of resorption when the WE formation was prevented.

**Figure 6:**
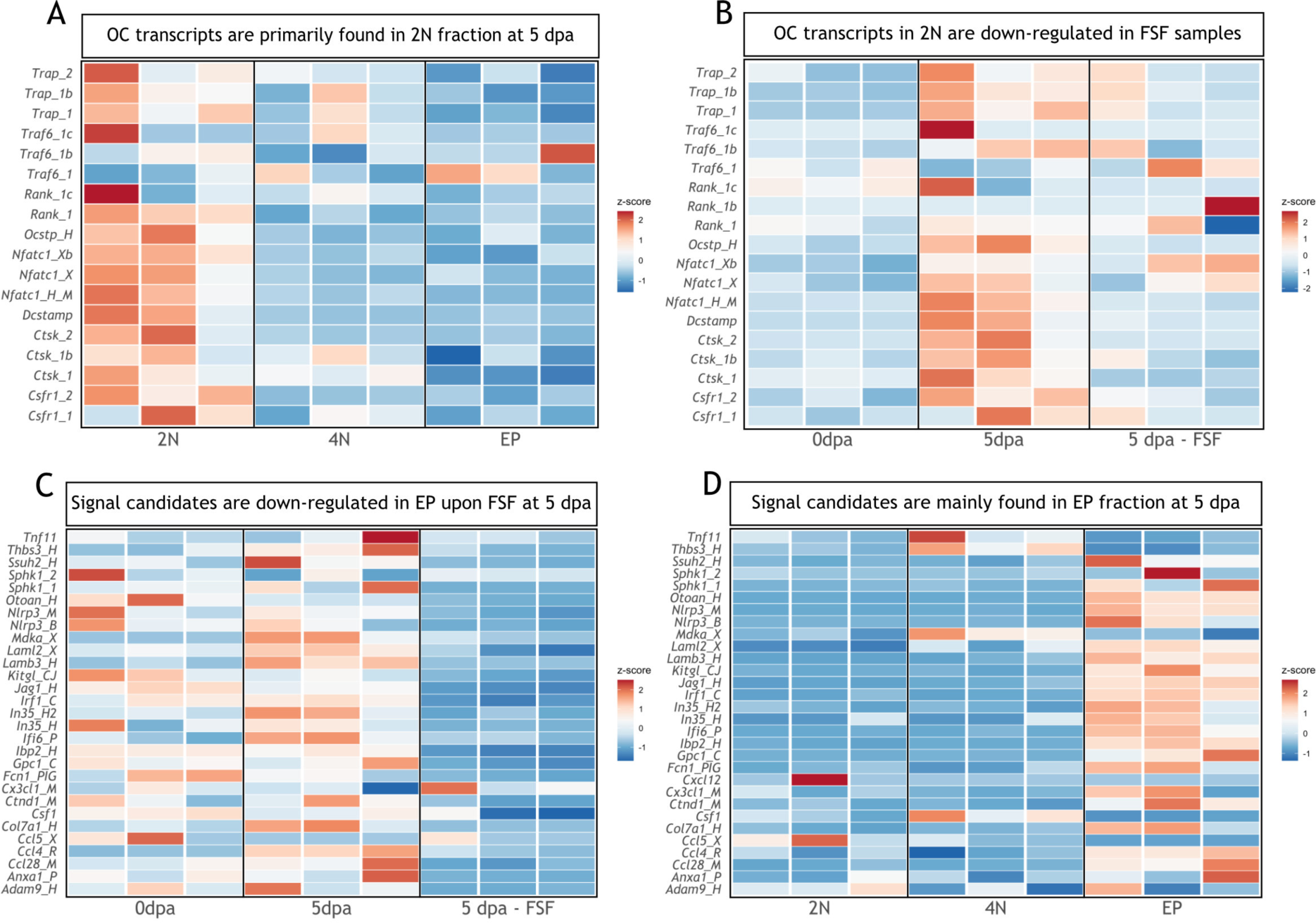
Transcripts associated with osteoclastogenesis are downregulated in FSF samples at 5 dpa. (A) Heatmap of transcripts associated with osteoclast function in three different fractions at 5 dpa (n = 3). 2N: mature cells, 4N: dividing cells, EP: epithelial cells. (B) Heatmap of transcripts associated with osteoclast function in 2N fraction at different time points (n = 3). FSF sample correspond at 5 dpa. (C) Heatmap of differentially down-regulated transcripts in EP fraction after FSF surgery at 5 dpa associated with osteoclast recruitment and/or differentiation (n = 3). (D) Heatmap of differentially down-regulated transcripts after FSF surgery in three different fractions associated with osteoclast recruitment and/or differentiation (n = 3).

Next, we evaluated which transcripts were significantly down-regulated in the EP fraction of FSF limbs compared to a control limb at 5 dpa (supplementary information Tsai et al., 2020). We found several transcripts associated with osteoclast recruitment and/or differentiation (Fig. 6C). From these candidates, previous work reports a role for *Ccl4* (Xuan et al., 2017), *Sphk1* (Baker et al., 2010; Ishii et al., 2009; Ryu et al., 2006) and *Mdka* (Maruyama et al., 2004) in osteoclastogenesis. Moreover, these three transcripts were up-regulated at 5 dpa compared to 0 dpa, suggesting their participation in regeneration (red arrows, Fig. 6C). Finally, we confirmed that most of the candidate transcripts shown in Fig. 6C were expressed in the EP fraction (Fig. 6D), including *Sphk1* and *Ccl4*. One of the exceptions was *Mdka*, whose levels were more prominent in the 4N fraction; however, it was recently shown that *Mdka* plays a critical role in WE development and inflammation control during the earlier stages of regeneration (Tsai et al., 2020). This analysis provides evidence that factors associated with osteoclastogenesis and osteoclast recruitment, normally produced by the WE, are down regulated when WE formation is inhibited. Correspondingly, osteoclast transcripts are down-regulated in blastema cells.

### Skeletal resorption and blastema formation are spatially and temporally correlated

Blastema formation is considered an accumulation of cells at the amputation plane. However, taking resorption into consideration, those cells are initially accumulating (or reprogramming) more proximal to the amputation plane. Here, we showed that resorption can reach up to 100% of the calcified tissue, and hence the accumulation of progenitor cells might occur up to 1 mm behind the amputation plane. With this in mind, we sought to assess the blastema specification in the context of skeletal resorption.

First, we measured the blastema surface in images taken at 15 dpa, when resorption is completed and blastema already formed in zol-treated animals. We considered the distal end of the skeletal elements as the starting point of blastema, as it was proposed that progenitor cells accumulate distal to the end of the skeleton (Tank et al., 1976). As shown in Fig. 7A, we found a significant decrease in blastema area in zol-treated animals (yellow dashed line). To efficiently analyze the blastema position during resorption, we used molecular markers. First, we performed whole mount EdU staining at different dpa (Fig. 7B). Similar to previous reports, in an intact limb, EdU^+^ cells are less than 0.5% of the total cells (Johnson et al., 2018). At 7 dpa, we observed EdU^+^ cells behind the amputation plane, spanning over more than 500 µm. These cells were located where we expected to observe resorption. Interestingly, several EdU^+^ cells were located in the periskeleton (arrowheads), which could account for cells contributing to skeletal regeneration (Currie et al., 2016; McCusker et al., 2016). These cells were found along most of the skeletal element length. At 10 dpa, when resorption is occurring, we observed a more defined blastema (white arrowhead), which contained EdU^+^ cells. Similar to 7 dpa, a significant proportion of those EdU^+^ cells were located next to the resorbing skeleton. Finally, at 15 dpa, we observed an evident reduction in the skeletal length (yellow arrowheads) and a defined blastema distal to those skeletal elements (white dashed line). At this point, very few EdU^+^ cells were found next to the skeleton.

**Figure 7:**
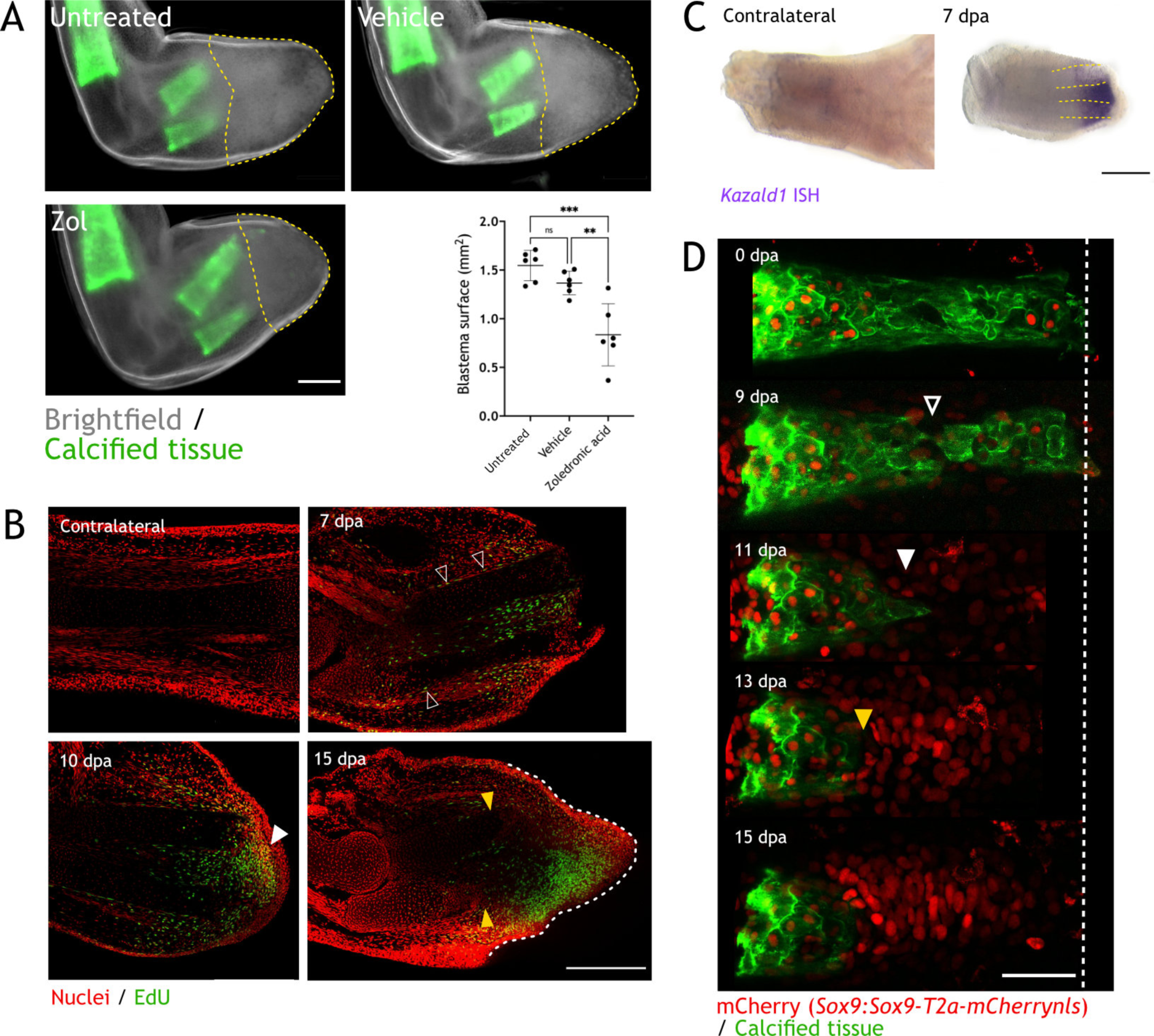
Skeletal resorption and blastema formation are spatially and temporally correlated. (A) Quantification of blastema size in zol treated limbs at 15 dpa. Dashed lines: blastema. Each dot represents an animal (n = 6, *** p < 0.001, ** p < 0.01, Tukey’s multiple comparisons test). (B) Whole mount EdU staining (green) in limbs upon zeugopod amputation at different dpa (n = 6). TO-PRO-3 was used for nuclear staining (red). Black arrowheads: dividing periskeletal cells. White arrowhead: blastema. Yellow arrowheads: distal end of skeletal element. Dashed line: blastema. Scale bar: 500 µm. (C) WISH for *Kazald1* in limbs upon zeugopod amputation at 7 dpa (n = 2). Yellow dashed lines: skeletal elements position. Scale bar: 500 µm. (D) Time course of resorption during digit regeneration in *Sox9-mCherry* (red) (n = 6). Calcein-stained (green) axolotls were amputated at the distal end of the calcified tissue. Black arrowhead: calcified tissue break. White arrowhead: condensation of mCherry*^+^* cells. Yellow arrowhead: resorption. Scale bar: 100 µm.

Although proliferation is mainly observed where blastema is forming, distal migration of EdU^+^ after division is also expected. The site of blastema formation was further assessed using a blastema marker in intact and 7 dpa limbs. We chose *Kazald1*, since it was shown to play a critical role in blastema formation (Bryant et al., 2017). Similar to the EdU^+^ labeling, we observed that *Kazald1*^+^ cells were located behind the amputation plane, surrounding the distal ends of the skeletal elements (dashed lines), in a zone where resorption is very likely to occur (Fig. 7C). These results suggest that blastema formation occurs in the same region and at the same time as resorption, and that using the skeletal element as a boundary for blastema identity, may provide an incomplete view of the course of regeneration.

Finally, to assess the cellular dynamics of skeletal resorption/formation, we used a *Sox9:Sox9- T2a-mCherrynls* transgenic line in conjunction with calcein staining to follow both processes *in vivo*. We performed amputation of the distal phalanx and followed the same amputated digit at different dpa (Fig. 7D). At 0 dpa, no mCherry^+^ cells were found outside the calcified tissue. At 9 dpa, we observed a break in the calcified phalanx (arrowhead) which shows a place where resorption was initiated. At 11 dpa, we observed a disorganized aggrupation of mCherry^+^ cells distal to the resorbed calcified tissue (white arrowhead). Those cells might represent the initial condensation of the regenerating cartilage. Interestingly, at 13 dpa, resorption continued (yellow arrowhead) and we saw an increase in the number and density of mCherry^+^ cells. Finally, at 15 dpa, resorption was finished and the condensation of mCherry^+^ cells in the new phalanx presented defined pattern. The condensation observed from 11 dpa, occurred behind the amputation plane, supporting our previous results. Moreover, these results show that resorption and skeletal regeneration are overlapping processes.

## DISCUSSION

Axolotl limb regeneration is an intricated multi-step process that requires the fine tuning of events such as wound closure, tissue repair, progenitor cells recruitment and the re-establishment of the functional unit. Although extensive work has been done to understand the cellular dynamics of blastema formation, other processes such as tissue histolysis, the immune response and tissue integration have yet to be fully understood.

Here, we report that upon digit and zeugopod amputation, significant skeletal resorption occurs which is carried out by an osteoclast population and can result in the resorption of 100% of the calcified matrix. Skeletal resorption is observed in young animals with cartilaginous limbs as well as young adult animals with ossified bones. Upon inhibition of resorption, we observed a clear failure in the integration of the regenerated zeugopodial elements and, interestingly, this failure can also occur in untreated animals undergoing regeneration. Moreover, we present strong evidence supporting the role of the WE in resorption induction. Finally, we observed a spatial correlation between resorption progression and blastema formation. Particularly, we found that the condensation of new cartilage started before the resorptive process is completed.

Similar to salamanders, mouse digit tip regeneration progresses with early histolysis and blastema formation. An extensive resorption of the phalanx can lead to bone volume reduction of almost 50% of its original size (Fernando et al., 2011). When osteoclasts are inhibited, regeneration is not compromised (Simkin et al., 2017) and when wound closure is induced earlier, resorption is delayed and regeneration progresses (Simkin et al., 2015). This suggest than in mouse digit tip resorption facilitates blastema formation. Moreover, it has been shown that both periosteal and endosteal cells are responsible for regeneration of the phalanx upon amputation in mice (Dawson et al., 2018), suggesting that bone resorption may be required for mobilization of a pool of progenitor cells. Although bone resorption is not required for regeneration to progress, this histolytic process is indeed an important event for the efficient regeneration of the mouse digit tip. In contrast to mouse, axolotl skeletal elements do not mobilize progenitors to the blastema and wound closure occurs within hours upon amputation, even when skeletal structures protrude at the surface. An important similarity to our findings in axolotl is the time frame when resorption occurs (7-15 dpa). This suggest that the unique fast activation and clearance of osteoclasts is particular to regeneration.

### Resorption efficiency defines skeletal integration success

In this work, osteoclast inhibition resulted in a clear failure in tissue integration. This phenotype often presented as an angulation of the radius, heterotopic cartilage formation in the ulna, or a complete separation between the mature and the regenerated structures. In general, we observed a higher rate of resorption for the ulna than for the radius, which could account for the more striking integration phenotypes reported for the radius.

Our experiments revealed a gradient of integration phenotypes correlated with the amount of tissue resorbed: the more resorption, the better the integration. Strikingly, in animals undergoing normal regeneration we often observed faulty integration phenotypes in mineralized skeleton, as seen by angulations at the stump-regenerated interphase. A recent report, showing abnormally regenerated limbs in almost 50% of the animals, showed some similar phenotypes to the ones presented here, i.e. a narrowing in the diaphysis and some heterotopic cartilage formation (Bothe et al., 2020). Using polarization microscopy, we presented evidence of an ECM disorganization in the interphase region where both stump and regenerated tissue are connected. Finally, we showed a prevalence of these phenotypes at 90 dpa, proving that they are not resolved after regeneration has been completed.

Remarkably, we observed a high variability in the amount of calcified tissue resorbed in different animals, ranging from 25 to 100% for radius and ulna, being the inter-assay variability higher than the intra-assay. The source of this variability could be an environmental factor (e.g. water temperature), but it highlights the different outcomes that skeletal regeneration can produce. Indeed, in some cases, resorption involved a sequential degradation of the skeletal tissue, and at other times, it involved the break and shedding of a skeleton piece, which has been observed in mouse digit tip (Fernando et al., 2011). In contrast to mouse, skeletal shedding in the axolotl is not associated with wound re-epithelialization, as this occurs after wound closure.

Our results highlight the misconception that axolotl limb regeneration recapitulates the pre-existing morphology with high fidelity. This report reveals that a faulty skeletal regeneration is a rather common outcome in the axolotl limb, and correlates resorption efficiency with successful skeletal integration.

### Wound epithelium, resorption and blastema

The WE is a critical structure for the progression of regeneration, regulating process such as ECM remodeling, tissue histolysis, proliferation and inflammation (Tsai et al., 2020). Here, by blocking the formation of the WE, we underscore its role in the initiation of resorption. Moreover, we analyzed available WE RNA-seq data and found several factors known to influence osteoclast progenitor migration and/or differentiation. Among those candidates, *Sphk1*, *Ccl4* and *Mdka* are up-regulated in the epithelial fraction during regeneration and have been linked to osteoclast biology. Indeed, S1P, which is phosphorylated by the sphingosine kinase (SPHK), has been shown to have a role in both bone resorption and bone formation (Ishii et al., 2009; Pederson et al., 2008; Ryu et al., 2006), while CCL4 and MDKA have been connected with osteoclast progenitors recruitment (Maruyama et al., 2004; Xuan et al., 2017). Future studies to understand how these factors are regulated in osteoclast-mediated resorption during regeneration will be needed.

We hypothesized that the connection between the WE and skeletal resorption could be mediated by a zone of influence determined by the WE position. Our results show that resorption starts distally, enclosed by the WE boundaries, supporting this idea. Although the WE may not be the only source of factors inducing osteoclast progenitor migration and differentiation, it probably is a general source of chemokines inducing the recruitment of RANK^+^ myeloid progenitors. We hypothesize that those myeloid cells will then recognize factors secreted by the skeletal elements that promote osteoclast differentiation. Indeed, the main source of RANKL in mammals are both hypertrophic chondrocytes and osteocytes (Xiong et al., 2011).

In this work, we also provide evidence supporting a spatiotemporal correlation between skeletal resorption and blastema formation. The WE produces signals involved in blastema proliferation and patterning (Boilly and Albert, 1990; Ghosh et al., 2008; Han et al., 2001; Tsai et al., 2020), and those signals could be acting in the same zone of influence. Moreover, since the skeletal elements are structural supports of the limb, the resorption of the hard matrix might cause a collapse of the soft tissue towards the proximal region, and thus a formation of the blastema behind the amputation plane. Indeed, we observed EdU^+^ cells and *Kazald1^+^* cells in the surroundings of the skeletal elements shortly before the start of resorption, and condensation of skeletal progenitors distal to the resorbed tissue and under the amputation plane.

Historically, the amputation plane has been conceptualized as a fixed position in the limb which determines the beginning of the blastema. However, the majority of the amputation experiments are performed by trimming the skeletal elements because this ensures consistency on the formation of a blastema. In this surgical procedure, the skin of the amputated limb is retracted and the extending skeletal elements are re-amputated. Trimming results in a faster WE and blastema formation; however, this procedure might cause the erroneous perception of a fixed amputation plane. Comparatively, in the case of the mouse digit tip, in which bone resorption occurs, a regeneration plane has been identified as more proximal than the amputation plane. (Seifert and Muneoka, 2018). This study has implications for demarcating the blastema, the mature cell source and the dynamic interphase created by nascent, migrating cells and a stream of morphogens.

### Future perspectives

There are still unresolved questions regarding the osteoclast population here described. First, what is the origin of this population during regeneration? Upon amputation, a peak of myeloid chemotactic molecules was reported at 1 dpa, followed by an infiltration of myeloid cells and macrophages (Godwin et al., 2013). This could suggest that osteoclast progenitors are recruited to the amputation plane, but it does not rule out the participation of resident progenitors in the neighboring tissues. Second, what is the fate of osteoclasts after resorption? We show here that this population eventually disappears and even shows some signs of apoptosis, but whether all undergo apoptosis or if some will recirculate, remains unknown. Current understanding of osteoclast biology suggest that they are short-lived cells, which undergo apoptosis after ca. 2 weeks (Manolagas, 2000). However, recent works have shown that osteoclasts can be longer lived (Jacome-Galarza et al., 2019) or be recycled via a cell type named osteomorphs (McDonald et al., 2021). These studies underline the need to continue investigating osteoclast biology *in vivo,* and particularly in this rapidly-triggered response to regeneration. The axolotl limb presents a new paradigm in which osteoclast function can be assessed, and thus the development of new transgenic lines to label myeloid progenitors and to indelibly label osteoclast will provide a mean to resolve the aforementioned questions.

In addition, the concomitant resorption and regeneration need to be further explored. It is known that histolysis helps with the mobilization of progenitor cells in salamanders (Thornton, 1938b) and in mouse (Dawson et al., 2018), but how osteoclast-mediated resorption could be influencing cartilage condensation in the context of axolotl regeneration remains to be studied. Specifically, how the cell differentiation and migration is orchestrated with respect to resorption is unclear. Of particular interest are periskeletal cells migrating from the periphery of the skeletal element towards the blastema, and contributing to the formation of the proximal skeleton. The cell source zone, i.e. a zone where blastema cells are recruited, has been roughly defined to be 500 µm from the amputation plane. However, resorption was not analyzed in that study (Currie et al., 2016). It is unclear how resorption and the detachment and migration of periskeletal cells are coordinated, or if the source of periskeletal cells correspond to a region not resorbed (e.g. proximal to the calcified tissue). Previous works have demonstrated the interaction between osteoclast and osteoblasts *in vivo* (Furuya et al., 2018; Ikebuchi et al., 2018), and the *in vivo* assessment of this in the context of skeletal regeneration would be necessary.

Finally, we need to consider the context in which resorption is occurring since different cell types are found in the same skeletal element along the proximo-distal axis, and this could influence the outcome of resorption in skeletal regeneration (Riquelme-Guzmán et al., 2021).

### Concluding remarks

This work presents a systematic assessment of the timing, extent and consequences of skeletal resorption. We show that the skeleton undergoes a massive and rapid histolytic event which is essential for a successful integration of the regenerated structure. This process, which is carried out by osteoclasts, is influenced by the formation of the WE and is correlated with the spatial position of the early blastema. Furthermore, we present proof that the axolotl limb regeneration is not perfect and it often leads to abnormal skeletal phenotypes. We consider that resorption is playing a key role in skeletal regeneration and its implications for regeneration needs to be further explored, particularly its coordination with cell migration and condensation of the new skeleton.

## MATERIALS AND METHODS

### Axolotl husbandry and transgenesis

Axolotls (*Ambystoma mexicanum*) were maintained at the CRTD axolotl facility and at Harvard University. All procedures were performed according to the Animal Ethics Committee of the State of Saxony, Germany, and the Institutional Animal Care and Use Committee (IACUC) Guidelines at Harvard University (Protocol 11-32). Animals used were selected by its size (snout to tail = ST). Most experiments were done using animals 4-6 cm ST, unless stated otherwise. We performed experiments using white axolotls (*d/d*). In addition, we utilized transgenic lines shown in table I.

**Table I:**
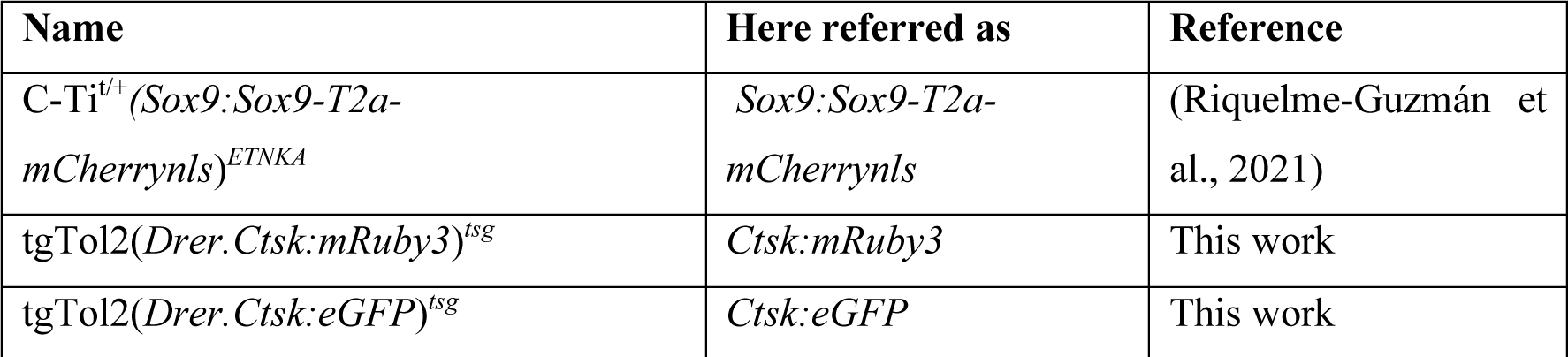
Transgenic lines used in this work.

To generate the *Ctsk:mRuby3* or *Ctsk:eGFP* transgenic lines, a plasmid containing 4 kb of *Ctsk* promoter from zebrafish together with *Tol2* sequences was used (kind gift from Knopf Lab at CRTD and Gilbert Weidinger at Ulm University). The *mRuby3* or *eGFP* coding region was cloned 3’ from the promoter. For ligation, plasmid restriction was performed using the FseI and XhoI restriction enzymes (#R0588S, #R0146S respectively; New England BioLabs, Frankfurt am Main, Germany). Fertilized embryos from *d/d* axolotls were injected with the *Ctsk:mRuby3* or *Ctsk:eGFP* vector and *Tol2* mRNA as previously described (Khattak et al., 2014). F0 animals were selected and grown in our colony until sexual maturity. For experiments, F0 were crossed with a *d/d* axolotl, and F1 animals were used.

### Experimental procedures in axolotls

*In vivo* skeletal staining was performed using calcein (#C0875, Sigma-Aldrich, Darmstadt, Germany) or alizarin red (#A5533, Sigma-Aldrich) A 0.1% solution was prepared for either dye with swimming water. Axolotls were submerged in solution for 5 - 10 minutes in the dark. After staining, animals were transferred to a tank with clean swimming water, which was changed as many times until water was not stained. Amputations were performed either 10 minutes after staining or the next day for better visualization.

For amputations, animals were anesthetized with 0.01% benzocaine solution. All amputations were performed at the distal end of the calcified diaphysis using an Olympus SZX16 stereomicroscope. After surgical procedure, animals were covered with a wet tissue (with benzocaine) and allowed to recover for 10 minutes prior to be transferred back to swimming water. The full skin flat surgery (FSF) was performed as described in (Tsai, 2020; Tsai et al., 2020).

Zoledronic acid (#SML0223, Sigma-Aldrich) treatment and EdU (#C10337, Invitrogen, Darmstadt, Germany) labelling were done by intraperitoneal injections in anesthetized axolotls. 200 µg/kg of zoledronic acid were injected every 3 days (stock 1 mg/mL in APBS (80% PBS)). 10 µg/g of EdU were injected 4 hours prior to tissue collection (stock 2.5 mg/mL in DMSO). Injection volume was adjusted to 10 µL with APBS. After injections, animals were kept covered with a wet paper for 10 minutes before returning them into the water tank.

*In vivo* imaging was performed in anesthetized animals. For stereoscope imaging, animals were placed in a 100 mm petri dish and limb was positioned accordingly. An Olympus SZX16 stereoscope microscope (objective: SDF Plapo 1xPF) was used. For confocal imaging, animals were place in a glass bottom dish (ø: 50/40 mm, #HBSB-5040, Willco Wells, Amsterdam, The Netherlands). A wet tissue with benzocaine was laid on top of the animal to avoid it to dry, and a silica block was laid on top of the hand to flatten it and improve light penetrance. A Zeiss confocal laser scanning microscope LSM780 (objectives: Plan apochromat 10x/0.45 or Plan-apochromat 20x/0.8) was used.

For tissue collection, animals were anesthetized prior to collection. After it, animals were euthanized by exposing them to lethal anesthesia (0.1% benzocaine) for at least 20 min. Tissue fixation and further procedures are described specifically for each case.

### Paraffin sectioning and Masson’s trichrome staining

Limbs were isolated and fixed with MEMFa 1x (MOPS 0.1M pH 7.4 / EGTA 2 mM / MgSO4x7H2O 1 mM / 3.7% formaldehyde) overnight at 4°C. Samples were washed with PBS and dehydrated with serial EtOH washes (25, 50, 70 and x3 100%). Samples were then incubated three times with Roti®Histol (#6640, Carl Roth, Karlsruhe, Germany) at RT and four times with paraffin (Roti®- Plast, #6642, Carl Roth) at 65°C in glass containers. After last incubation, samples were embedded in paraffin using plastic containers and stored at RT. Longitudinal sections of 6 µM thickness were obtained.

Masson’s trichrome staining on paraffin sections was performed following the producer’s recommendations (Procedure No. HT15, Sigma-Aldrich). Imaging was performed in a Zeiss Axio Observer.Z1 inverted microscope (objective: Plan-apochromat 20x/0.8).

### Cryosectioning

Limbs fixed with MEMFa were washed with PBS and decalcified with EDTA 0.5 M at 4°C for 48 hours. Next, limbs were washed with PBS and incubated overnight with sucrose 30% at 4°C. Samples were embedded in O.C.T. compound (#4583, Tissue-Tek, Umkirch, Germany) using plastic molds and frozen with dry ice for 1 hour prior to storage at -20°C. Longitudinal sections of 12 µm thickness were cut and mounted on superfrost slides. Slides were kept at -20°C until processed.

### TRAP enzymatic staining

Tartrate-resistant acid phosphatase (TRAP) enzymatic staining was performed in cryosections. Slides were dried for 1 hour prior to wash them with PBS + 0.1% Tween-20 for 10 minutes. Next, slides were permeabilised with PBS + 0.3% Tx-100 for 1 hour. After permeabilization, slides were equilibrated by three washes with TRAP buffer (NaAcetate 0.1M / acetic acid 0.1M / NaTartrate 50 mM / pH 5.2) for 10 minutes at 37°C in water bath. Slides were stained with color reaction buffer (TRAP buffer / Naphthol AS-MX phosphate 1.5 mM / Fast Red Violet LB Salt 0.5 mM) for 1 hour at 37°C in water bath. After staining, slides were washed three times with PBS for 10 minutes and mounted with Entellan^TM^ (#1.07960, Sigma-Aldrich). Images were taken in a Zeiss Axio Observer.Z1 inverted microscope.

### Immunofluorescence

For immunofluorescence (IF), cryosections were used. Slides were dried at RT for at least 1 hour. Sections were washed three times with PBS + 0.3% Tx-100 prior to blocking with PBS + 0.3% Tx-100 + 10% normal horse serum (NHS, #S-2000-20, Vector Labs, Burlingame, CA, USA) for 1 hour. Primary anti-CTSK (#ab19027, Abcam, Cambridge, UK) antibody incubation (1:20) was done in blocking solution for 1 hour at RT and then overnight at 4°C. Sections were then washed three times with PBS + 0.3% Tx-100 and incubated with Goat anti-Rabbit, Alexa Fluor 647 antibody (1:200, #A- 21245, Invitrogen) and Hoechst 33342 1:1000 for 2 hours. Finally, sections were washed three times with PBS + 0.3% Tx-100 and mounted using Mowiol mounting medium (#0713 Carl Roth). Imaging was performed on a Zeiss Axio Observer.Z1 inverted microscope with an ApoTome1 system (objectives: Plan-apochromat 10x/0.45 or Plan-apochromat 20x/0.8).

### RNA probes for *in situ* hybridization

*Ctsk, Kazald1* and *Krt17* probes were created by TA cloning. Probe amplification was done using primers previously published (Table II). Ligation was done into a pGEM®-T easy vector system I (#A1360, Promega, Madison, WI, USA). To confirm successful cloning, vectors were purified and sequenced using the Mix2Seq Kit (Eurofins Genomics, Ebersberg, Germany).

**Table II:**
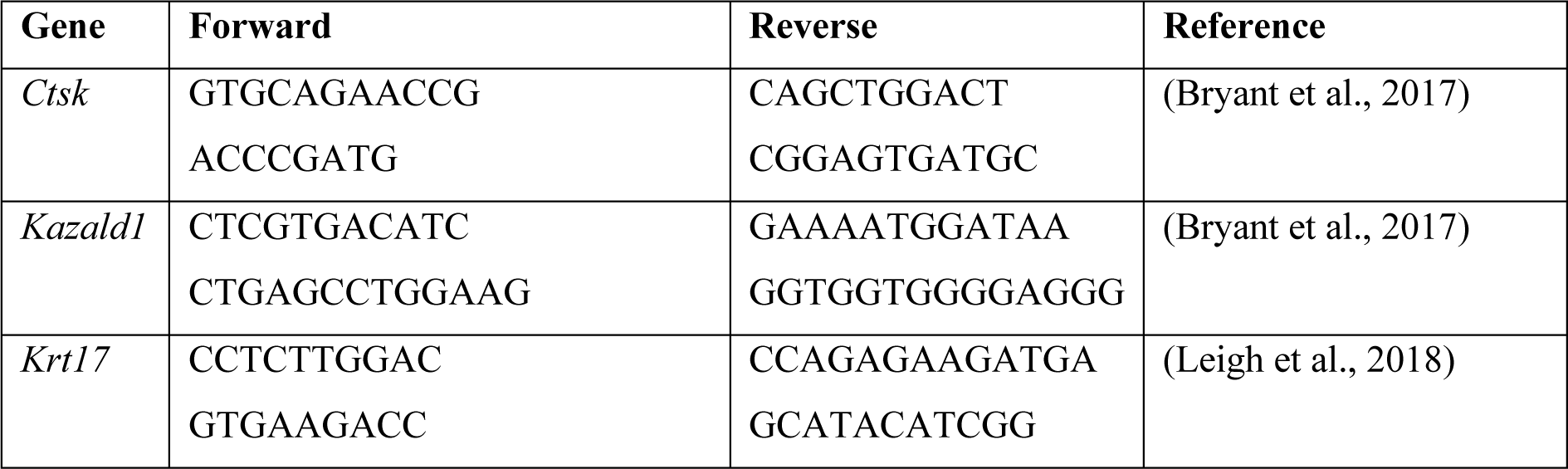
Primers for ISH probes cloning.

For synthesizing the ISH probes, *in vitro* transcription was carried out using a T7 polymerase (#RPOLT7-RO, Roche, Mannheim, Germany) or a SP6 polymerase (#RPOLSP6-RO, Roche) following provider’s instructions. Prior to transcription, 5 µg of plasmid were linearized using the restriction enzyme SpeI-HF® (#R3133S, New England BioLabs) for *Ctsk* and *Krt17*, or SphI-HF® (#R3182S, New England BioLabs) for *Kazald1*. Probes were purified using the RNAeasy® Mini Kit (#74104, QIAGEN, Hilden, Germany) according to provider’s instructions.

### *In situ* hybridization (ISH)

ISH was performed in cryosections using *Ctsk* probe following a previously published protocol (Knapp et al., 2013). When the protocol was finished, slides were fixed in formaldehyde 4% overnight at RT. Slides were then dehydrated with serial EtOH washes (25, 50, 70 and 100%) prior to wash with Roti®Histol and mounting with Entellan^TM^. Imaging was performed on a Zeiss Axio Observer.Z1 inverted microscope.

### Whole mount *in situ* hybridization (WISH)

Whole mount ISH was performed using *Krt17* and *Kazald1* probes. Protocol was adapted from (Woltering et al., 2009). Briefly, samples were dehydrated with serial MetOH washes (25, 50, 75% in PBS + 0.1% Tween-20 and 100%). Limbs were bleached in MetOH + 6% H2O2 at RT and then rehydrated with serial washes of MetOH. Then, limbs were washed with TBST (1x TBS, 0.1% Tween-1. 20) and treated with 20 µg/mL Proteinase K in TBST for 30 min at 37°C. After incubation, limbs were washed with TBST and rinsed with trietanolamine 0.1M pH 7.5 (#90278, Sigma-Aldrich). Limbs were incubated with freshly prepared 0.1M TEA + 1% acetic anhydride (#320102, Sigma-Aldrich) for 10 minutes and then washed again with TBST. Next, limbs were fixed with 4% PFA + 0.2% glutaraldehyde (#G6257, Sigma-Aldrich) for 20 minutes and washed with TBST. TBST was removed and limbs were incubated with previously warmed Pre-Hyb solution (hybridization buffer without probe) at 60 °C for 4 hours, prior to be transferred into pre-warmed hybridization buffer + probe (6 µL/mL) and incubated overnight at 60°C. The next day, limbs were washed at 60°C with pre-warmed 5xSSC solution twice for 30 minutes, with 2xSSC solution three times for 20 minutes, and with 0.2xSSC three times for 20 minutes. Limbs were then washed with TNE solution twice for 10 minutes at RT prior to incubation with 20 µg/mL RNAse A in TNE solution for 15 minutes. After incubation, limbs were washed with TNE solution twice for 10 minutes, and with MAB solution three times for 5 minutes. Limbs were blocked with MAB solution + 1% blocking reagent for 1.5 hours, and then incubated with MAB solution + 1% blocking reagent + 1:3000 anti-digoxigenin-AP, Fab fragments for 4 hours at RT. Next, limbs were washed with MAB solution three times for 5 minutes each and then overnight at RT. On day 3, limbs were washed with MAB solution five times for 1 hour each and again overnight. After MAB washes, limbs were washed with NTMT three times for 10 minutes at RT, and then incubated with freshly made NTMT + 20 µL/mL NBT/BCIP (#11681451001, Roche). For both *Kazald1* and *Krt17* probes, 4-6 hours of incubation were enough for signal to develop. Reaction was then stopped by incubating with PBS + 0.1% Tween-20 twice for 10 minutes and then fixing with 4% PFA at 4°C overnight. After fixation, limbs were washed with PBS + 0.1% Tween-20 and stored in that solution at RT. Imaging was performed on a Zeiss Discovery.V20 stereomicroscope.

### Alcian blue/alizarin red staining

Staining was performed as recently described (Riquelme-Guzmán et al., 2021). Imaging was performed on a Zeiss Discovery.V20 stereomicroscope (objective: Plan S 1.0x).

### Whole mount EdU staining

Limbs from axolotls injected with EdU were fixed with MEMFa 1x overnight at 4°C and then washed with PBS. For whole mount EdU staining, limbs were washed with PBS + 0.3% Tx-100 twice for 2 hours and then blocked with PBS + 0.3% Tx-100 + 5% goat serum + 10% DMSO for 24 hours at RT. Click-iT^TM^ EdU cell proliferation kit, Alexa Fluor 488 (#C10337, Invitrogen) was used following provider’s instructions. Samples were incubated in reaction cocktail for 4 hours at RT. After incubation, samples were washed with PBS + 0.3% Tx-100 four times for 15 min. Next, samples were incubated with TO-PRO^TM^-3 1:10.000 in PBS + 0.3% Tx-100 for 1 hour at RT. Finally, limbs were washed with PBS four times for 15 min each at RT. Limbs were cleared by dehydration with serial washes of EtOH (25, 50, 70, 100%) for 2 hours each at 4°C. Samples were then incubated overnight in 100% EtOH at 4°C prior to clearing with ethyl cinnamate (#112372, Sigma-Aldrich) at RT for at least 2 hours. Samples were imaged the same day on a Zeiss confocal laser scanning microscope LSM 780.

### RNA purification and RT-qPCR

Limbs for RNA isolation were stored in RNA*later*^TM^ (#AM7024, Invitrogen) at -20°C until all samples were collected. RNA isolation was performed using the RNAeasy® Mini Kit. 50 ng of RNA were used for cDNA synthesis using the PrimeScript^TM^ RT reagent Kit (#RR037A, Takara, Göteborg, Sweden) following the provider’s instructions. RT-qPCR was performed using the TB Green® Premix Ex Taq^TM^ (Tli RNAseH Plus) kit (#RR420A, Takara). RT-qPCR was done using a LightCycler 480 system with a pre-defined protocol for SYBR Green. Results were analyzed using the ΔΔCT method and the *Rpl4* housekeeping gene. After analysis, results were shown as relative levels compared to a control. Primer pairs used are listed in Table III.

**Table III:**
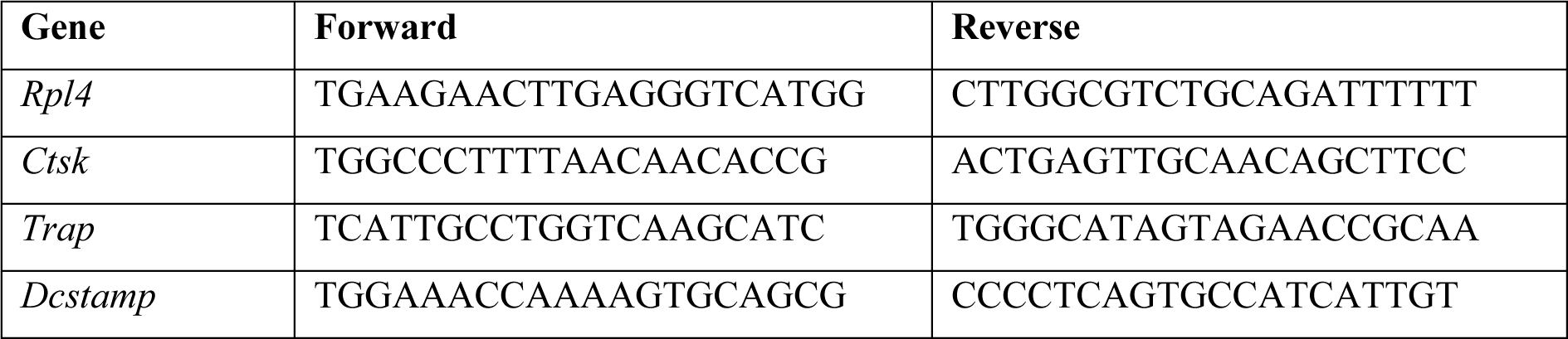
Primer pairs used for RT-qPCR.

### Polarization microscopy

The LC-PolScope is a powerful tool to quantitatively image optically anisotropic materials having a refractive index that depends on the polarization and propagation of light (birefringence), such as collagen, the main component of cartilage ECM (Fox et al., 2009). An LC-PolScope (on a Ti Eclipse microscope body) with a sCMOS camera (Hamamatsu Orca Flash 4.0) was used. Acquisitions were done with a 20x/0.8 objective and using µManager software (Edelstein et al., 2014). Two images were acquired: the retardance and the slow axis orientation. The retardance correlates with the amount of birefringent components, while the slow axis orientation image provides information on the orientation of those components, i.e. the angle in which they are aligned in the sample. Retardance and slow axis orientation images were aligned using a custom-made MATLAB script such that the x-axis corresponded to the proximodistal axis and the y-axis corresponded to the anteroposterior axis, with the elbow on the top-right corner of the image. The angle was measured respect to the proximodistal axis. Once the images were aligned, the regions of interest were cropped and segmented using the Trainable Weka Segmentation plugin from Fiji (Arganda-Carreras et al., 2017). The segmentation was done to obtain masks for the collagen regions and to remove the cells from the analysis. The masks were then applied to the slow axis orientation images and the orientations of the collagen fibers were quantified using MATLAB.

### Curating RNA-seq data

Recently published axolotl RNA-seq datasets (Tsai et al., 2020, 2019) were used to evaluate osteoclasts-related transcripts levels in samples upon full skin flap surgery. For curating the datasets, R Studio was used (RStudio Team, http://www.rstudio.com/). In order to find osteoclasts-related transcripts identifiers, the human, mouse and xenopus protein orthologous for each transcript were used. With the protein sequences, a protein blast was performed using the predicted proteins from Bryant et al. *de novo* axolotl transcriptome (supplementary data Bryant et al., 2017). The best three matches for each protein were used for browsing in Tsai’s transcriptome. The datasets from both of Tsai et al. works were combined and filtered in order to have only the results from 0 dpa, 5 dpa and 5 dpa in FSF surgery. In addition, the 2N, 4N and EP fractions at 5 dpa were also filtered from the combined datasets. For organizing, filtering and calculating z-scores in the datasets, the tidyverse package (Wickham et al., 2019) and plyr package (Wickham, 2011) were used. For creating the heatmaps, the ggplot2 package was used (Wickham, 2016).

To find possible candidates involved in osteoclast recruitment and differentiation, a search in the available literature was done for each differentially down-regulated transcript in FSF samples (385 transcripts, Tsai et al., 2020 supplementary data). Transcripts which have been connected to osteoclast function or belong to a protein family shown to play a role in osteoclast recruitment and differentiation, were filtered and heatmaps were created for better visualization of the levels during regeneration and in the different fractions (Fig 18C, D).

### µCT scan

Scans were performed as recently described (Riquelme-Guzmán et al., 2021). Threshold was set to 220 mg HA/cm^3^.

### Statistical analysis

Statistical analyses were performed using the software Prism9 (GraphPad Software, LLC, San Diego, CA, USA) for macOS. Statistical tests performed are described in each figure. P-values < 0.05 were considered statistically significant.

### Image processing and figure design

All images were processed using Fiji (Schindelin et al., 2012). Processing involved selecting regions of interest, merging or splitting channels and improving brightness levels for proper presentation in figures. Maximum intensity projections were done in some confocal images and it is stated in the respective figure’s descriptions. Stitching of tiles was done directly in the acquisition software Zen (Zeiss Microscopy, Jena, Germany). Figures were created using Affinity Designer (Serif Europe, West Bridgford, UK).

## ACKNOWLEDGMENTS

We thank all the members from the Sandoval-Guzmán lab for continuous support during the development of this work. We are also grateful to Anja Wagner, Beate Gruhl and Judith Konantz for their impeccable dedication to the axolotl care. This work was funded by a DFG Research Grant (432439166). CRG was supported by the DIGS-BB Fellow award. This work was supported by the Light Microscopy Facility, a Core Facility of the CMCB Technology Platform at TU Dresden.

## COMPETING INTERESTS

No conflicts of interest, financial or otherwise, are declared by the authors.

---

**Figure S1:**
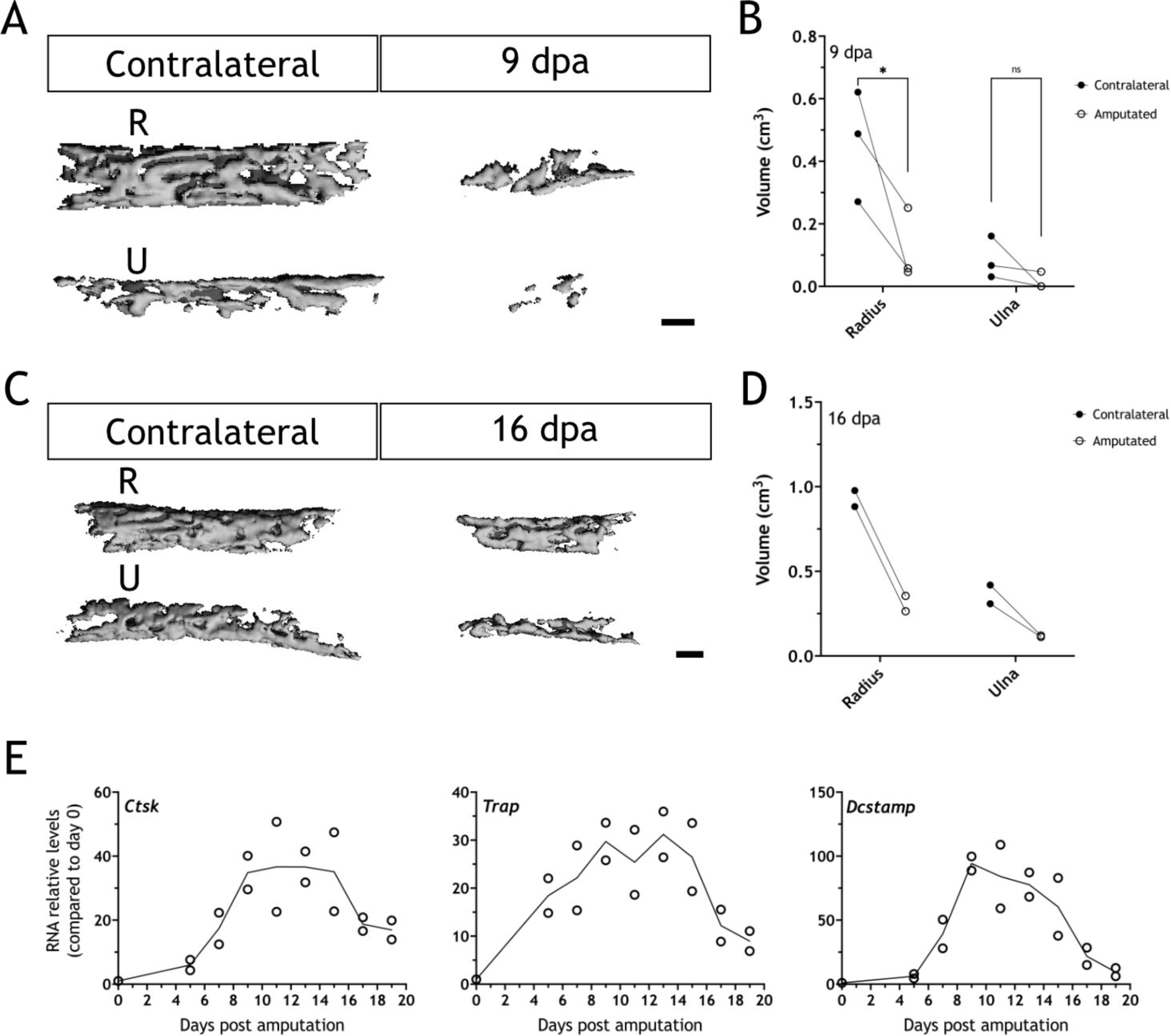
Bones are resorbed upon amputation in 16 cm ST axolotls. (A) 3D reconstructions from µCT scans for radius (R) and ulna (U) in the contralateral and a 9 dpa limb upon amputation at the distal end of the calcified tissue (n = 3). Scale bar: 200 µm. (B) Quantification of bones volume (cm^3^) for samples in (A). Each dot represents an animal (n = 3. * p < 0.05, Bonferroni’s multiple comparisons test, contralateral versus amputated). (C) 3D reconstructions from µCT scans for radius and ulna in the contralateral and a 16 dpa limb upon amputation at the distal end of the calcified tissue (n = 2). Scale bar: 200 µm. (D) Quantification of bones volume (cm^3^) for samples in (C). Each dot represents an animal (n = 2). (E) RT-qPCR for C*tsk, Trap* and D*cstamp* at different dpa upon zeugopodial amputation. Solid line represents mean, each dot is an animal (n = 2).

## Notes

### Competing Interest Statement

The authors have declared no competing interest.

## REFERENCES

Arganda-Carreras I, Kaynig V, Rueden C, Eliceiri KW, Schindelin J, Cardona A, Seung HS. 2017. Trainable Weka Segmentation: a machine learning tool for microscopy pixel classification. Bioinformatics 33:2424–2426. doi:10.1093/bioinformatics/btx180

Aztekin C. 2021. Tissues and Cell Types of Appendage Regeneration: A Detailed Look at the Wound Epidermis and Its Specialized Forms. Front Physiol 12:771040. doi:10.3389/fphys.2021.771040

Baker DA, Barth J, Chang R, Obeid LM, Gilkeson GS. 2010. Genetic Sphingosine Kinase 1 Deficiency Significantly Decreases Synovial Inflammation and Joint Erosions in Murine TNF-α– Induced Arthritis. J Immunol 185:2570–2579. doi:10.4049/jimmunol.1000644

Blum N, Begemann G. 2015. Osteoblast de- and redifferentiation are controlled by a dynamic response to retinoic acid during zebrafish fin regeneration. Development 142:2894–2903. doi:10.1242/dev.120204

Boilly B, Albert P. 1990. In vitro control of blastema cell proliferation by extracts from epidermal cap and mesenchyme of regenerating limbs of axolotls. Roux’s Archives Dev Biology 198:443–447. doi:10.1007/bf00399054

Bothe V, Mahlow K, Fröbisch NB. 2020. A histological study of normal and pathological limb regeneration in the Mexican axolotl Ambystoma mexicanum. J Exp Zoology Part B Mol Dev Evol. doi:10.1002/jez.b.22950

Bryant DM, Johnson K, DiTommaso T, Tickle T, Couger M, Payzin-Dogru D, Lee TJ, Leigh ND, Kuo T-H, Davis FG, Bateman J, Bryant S, Guzikowski AR, Tsai SL, Coyne S, Ye WW, Freeman RM, Peshkin L, Tabin CJ, Regev A, Haas BJ, Whited JL. 2017. A Tissue-Mapped Axolotl De Novo Transcriptome Enables Identification of Limb Regeneration Factors. Cell Reports 18:762– 776. doi:10.1016/j.celrep.2016.12.063

Cappariello A, Maurizi A, Veeriah V, Teti A. 2014. The Great Beauty of the osteoclast. Archives of Biochemistry and Biophysics 558:70–78. doi:10.1016/j.abb.2014.06.017

Charles JF, Aliprantis AO. 2014. Osteoclasts: more than ‘bone eaters.’ Trends in Molecular Medicine 20:449–459. doi:10.1016/j.molmed.2014.06.001

Clézardin P. 2013. Mechanisms of action of bisphosphonates in oncology: a scientific concept evolving from antiresorptive to anticancer activities. Bonekey Reports 2:267. doi:10.1038/bonekey.2013.1

Currie JD, Kawaguchi A, Traspas R, Schuez M, Chara O, Tanaka EM. 2016. Live Imaging of Axolotl Digit Regeneration Reveals Spatiotemporal Choreography of Diverse Connective Tissue Progenitor Pools. Dev Cell 39:411–423. doi:10.1016/j.devcel.2016.10.013

Dawson LA, Schanes PP, Kim P, Imholt FM, Qureshi O, Dolan CP, Yu L, Yan M, Zimmel KN, Falck AR, Muneoka K. 2018. Blastema formation and periosteal ossification in the regenerating adult mouse digit. Wound repair and regeneration : official publication of the Wound Healing Society [and] the European Tissue Repair Society 26:263–273. doi:10.1111/wrr.12666

Debuque RJ, Hart AJ, Johnson GH, Rosenthal NA, Godwin JW. 2021. Identification of the Adult Hematopoietic Liver as the Primary Reservoir for the Recruitment of Pro-regenerative Macrophages Required for Salamander Limb Regeneration. Frontiers Cell Dev Biology 9:750587. doi:10.3389/fcell.2021.750587

Dhillon S. 2016. Zoledronic Acid (Reclast®, Aclasta®): A Review in Osteoporosis. Drugs 76:1683– 1697. doi:10.1007/s40265-016-0662-4

Dunis DA, Namenwirth M. 1977. The role of grafted skin in the regeneration of X-irradiated axolotl limbs. Dev Biol 56:97–109. doi:10.1016/0012-1606(77)90157-9

Edelstein AD, Tsuchida MA, Amodaj N, Pinkard H, Vale RD, Stuurman N. 2014. Advanced methods of microscope control using μManager software. J Biological Methods 1:e10. doi:10.14440/jbm.2014.36

Einhorn TA, Gerstenfeld LC. 2015. Fracture healing: mechanisms and interventions. Nat Rev Rheumatol 11:45–54. doi:10.1038/nrrheum.2014.164

Fernando WA, Leininger E, Simkin J, Li N, Malcom CA, Sathyamoorthi S, Han M, Muneoka K. 2011. Wound healing and blastema formation in regenerating digit tips of adult mice. Developmental biology 350:301–10. doi:10.1016/j.ydbio.2010.11.035

Fischman DA, Hay ED. 1962. Origin of osteoclasts from mononuclear leucocytes in regenerating newt limbs. Anatomical Rec 143:329–337. doi:10.1002/ar.1091430402

Fox AJS, Bedi A, Rodeo SA. 2009. The Basic Science of Articular Cartilage. Sports Heal 1:461–468. doi:10.1177/1941738109350438

Furuya M, Kikuta J, Fujimori S, Seno S, Maeda H, Shirazaki M, Uenaka M, Mizuno H, Iwamoto Y, Morimoto A, Hashimoto K, Ito T, Isogai Y, Kashii M, Kaito T, Ohba S, Chung U, Lichtler AC, Kikuchi K, Matsuda H, Yoshikawa H, Ishii M. 2018. Direct cell–cell contact between mature osteoblasts and osteoclasts dynamically controls their functions in vivo. Nature Communications 9:300. doi:10.1038/s41467-017-02541-w

Gerber T, Murawala P, Knapp D, Masselink W, Schuez M, Hermann S, Gac-Santel M, Nowoshilow S, Kageyama J, Khattak S, Currie J, Camp GJ, Tanaka EM, Treutlein B. 2018. Single-cell analysis uncovers convergence of cell identities during axolotl limb regeneration. Science eaaq0681. doi:10.1126/science.aaq0681

Ghosh S, Roy S, Séguin C, Bryant SV, Gardiner DM. 2008. Analysis of the expression and function of Wnt-5a and Wnt-5b in developing and regenerating axolotl (Ambystoma mexicanum) limbs. Dev Growth Differ 50:289–297. doi:10.1111/j.1440-169x.2008.01000.x

Godwin JW, Pinto AR, Rosenthal NA. 2013. Macrophages are required for adult salamander limb regeneration. Proceedings of the National Academy of Sciences of the United States of America 110:9415–20. doi:10.1073/pnas.1300290110

Han M, An J, Kim W. 2001. Expression patterns of Fgf-8 during development and limb regeneration of the axolotl. Dev Dyn 220:40–48. doi:10.1002/1097-0177(2000)9999:9999<::aid- dvdy1085>3.0.co;2-8

Hay ED, Fischman DA. 1961. Origin of the blastema in regenerating limbs of the newt Triturus viridescens: An autoradiographic study using tritiated thymidine to follow cell proliferation and migration. Developmental Biology 3:26–59. doi:https://doi.org/10.1016/0012-1606(61)90009-4

Huang T, Zuo L, Walczyńska KS, Zhu M, Liang Y. 2021. Essential roles of matrix metalloproteinases in axolotl digit regeneration. Cell Tissue Res 385:105–113. doi:10.1007/s00441-021-03434-7

Hutchison C, Pilote M, Roy S. 2007. The axolotl limb: A model for bone development, regeneration and fracture healing. Bone 40:45–56. doi:10.1016/j.bone.2006.07.005

Ikebuchi Y, Aoki S, Honma M, Hayashi M, Sugamori Y, Khan M, Kariya Y, Kato G, Tabata Y, Penninger JM, Udagawa N, Aoki K, Suzuki H. 2018. Coupling of bone resorption and formation by RANKL reverse signalling. Nature 561:195–200. doi:10.1038/s41586-018-0482-7

Ishii M, Egen JG, Klauschen F, Meier-Schellersheim M, Saeki Y, Vacher J, Proia RL, Germain RN. 2009. Sphingosine-1-phosphate mobilizes osteoclast precursors and regulates bone homeostasis. Nature 458:524. doi:10.1038/nature07713

Jacome-Galarza CE, Percin GI, Muller JT, Mass E, Lazarov T, Eitler J, Rauner M, Yadav VK, Crozet L, Bohm M, Loyher P-L, Karsenty G, Waskow C, Geissmann F. 2019. Developmental origin, functional maintenance and genetic rescue of osteoclasts. Nature 1–5. doi:10.1038/s41586-019-1105-7

Johnson K, Bateman J, DiTommaso T, Wong AY, Whited JL. 2018. Systemic cell cycle activation is induced following complex tissue injury in axolotl. Developmental Biology 433:461–472. doi:10.1016/j.ydbio.2017.07.010

Khattak S, Murawala P, Andreas H, Kappert V, Schuez M, Sandoval-Guzmán T, Crawford K, Tanaka EM. 2014. Optimized axolotl (Ambystoma mexicanum) husbandry, breeding, metamorphosis, transgenesis and tamoxifen-mediated recombination. Nat Protoc 9:529–540. doi:10.1038/nprot.2014.040

Knapp D, Schulz H, Rascon C, Volkmer M, Scholz J, Nacu E, Le M, Novozhilov S, Tazaki A, Protze S, Jacob T, Hubner N, Habermann B, Tanaka EM. 2013. Comparative Transcriptional Profiling of the Axolotl Limb Identifies a Tripartite Regeneration-Specific Gene Program. PLoS ONE 8:e61352. doi:10.1371/journal.pone.0061352

Kozhemyakina E, Lassar AB, Zelzer E. 2015. A pathway to bone: signaling molecules and transcription factors involved in chondrocyte development and maturation. Development 142:817– 31. doi:10.1242/dev.105536

Kragl M, Knapp D, Nacu E, Khattak S, Maden M, Epperlein H, Tanaka EM. 2009. Cells keep a memory of their tissue origin during axolotl limb regeneration. Nature 460:60–65. doi:10.1038/nature08152

Leigh ND, Dunlap GS, Johnson K, Mariano R, Oshiro R, Wong AY, Bryant DM, Miller BM, Ratner A, Chen A, Ye WW, Haas BJ, Whited JL. 2018. Transcriptomic landscape of the blastema niche in regenerating adult axolotl limbs at single-cell resolution. Nature Communications 9:5153. doi:10.1038/s41467-018-07604-0

Manolagas SC. 2000. Birth and Death of Bone Cells: Basic Regulatory Mechanisms and Implications for the Pathogenesis and Treatment of Osteoporosis. Endocr Rev 21:115–137. doi:10.1210/edrv.21.2.0395

Maruyama K, Muramatsu H, Ishiguro N, Muramatsu T. 2004. Midkine, a heparin-binding growth factor, is fundamentally involved in the pathogenesis of rheumatoid arthritis. Arthritis Rheumatism 50:1420–1429. doi:10.1002/art.20175

McCusker CD, Diaz-Castillo C, Sosnik J, Phan AQ, Gardiner DM. 2016. Cartilage and bone cells do not participate in skeletal regeneration in Ambystoma mexicanum limbs. Dev Biol 416:26–33. doi:10.1016/j.ydbio.2016.05.032

McDonald MM, Khoo WH, Ng PY, Xiao Y, Zamerli J, Thatcher P, Kyaw W, Pathmanandavel K, Grootveld AK, Moran I, Butt D, Nguyen A, Corr A, Warren S, Biro M, Butterfield NC, Guilfoyle SE, Komla-Ebri D, Dack MRG, Dewhurst HF, Logan JG, Li Y, Mohanty ST, Byrne N, Terry RL, Simic MK, Chai R, Quinn JMW, Youlten SE, Pettitt JA, Abi-Hanna D, Jain R, Weninger W, Lundberg M, Sun S, Ebetino FH, Timpson P, Lee WM, Baldock PA, Rogers MJ, Brink R, Williams GR, Bassett JHD, Kemp JP, Pavlos NJ, Croucher PI, Phan TG. 2021. Osteoclasts recycle via osteomorphs during RANKL-stimulated bone resorption. Cell 184:1330–1347.e13. doi:10.1016/j.cell.2021.02.002

Mescher AL. 1976. Effects on adult newt limb regeneration of partial and complete skin flaps over the amputation surface. J Exp Zool 195:117–127. doi:10.1002/jez.1401950111

Muneoka K, Fox WF, Bryant SV. 1986. Cellular contribution from dermis and cartilage to the regenerating limb blastema in axolotls. Developmental Biology 116:256–260. doi:10.1016/0012-1606(86)90062-X

Neufeld DA, Aulthouse AL. 1986. Association of mesenchyme with attenuated basement membranes during morphogenetic stages of newt limb regeneration. The American Journal of Anatomy 176. doi:https://doi.org/10.1002/aja.1001760404

Nguyen M, Singhal P, Piet J, Shefelbine SJ, Maden M, Voss RS, Monaghan JR. 2017. Retinoic acid receptor regulation of epimorphic and homeostatic regeneration in the axolotl. Development 144:dev.139873. doi:10.1242/dev.139873

Oldenbourg R. 1996. A new view on polarization microscopy. Nature 381:811–812. doi:10.1038/381811a0

Pederson L, Ruan M, Westendorf JJ, Khosla S, Oursler MJ. 2008. Regulation of bone formation by osteoclasts involves Wnt/BMP signaling and the chemokine sphingosine-1-phosphate. Proc National Acad Sci 105:20764–20769. doi:10.1073/pnas.0805133106

Repesh LA, Oberpriller JC. 1978. Scanning electron microscopy of epidermal cell migration in wound healing during limb regeneration in the adult newt, Notophthalmus viridescens. Am j anat 151:539–555. doi:https://doi.org/10.1002/aja.1001510408

Riquelme-Guzmán C, Schuez M, Böhm A, Knapp D, Edwards-Jorquera S, Ceccarelli AS, Chara O, Rauner M, Sandoval-Guzmán T. 2021. Postembryonic development and aging of the appendicular skeleton in Ambystoma mexicanum. Dev Dynam. doi:10.1002/dvdy.407

Rodgers AK, Smith JJ, Voss SR. 2020. Identification of immune and non-immune cells in regenerating axolotl limbs by single-cell sequencing. Exp Cell Res 394:112149. doi:10.1016/j.yexcr.2020.112149

Ryu J, Kim HJ, Chang E, Huang H, Banno Y, Kim H. 2006. Sphingosine 1-phosphate as a regulator of osteoclast differentiation and osteoclast–osteoblast coupling. Embo J 25:5840–5851. doi:10.1038/sj.emboj.7601430

Sandoval-Guzmán T, Currie JD. 2018. The journey of cells through regeneration. Current opinion in cell biology 55:36–41. doi:10.1016/j.ceb.2018.05.008

Schindelin J, Arganda-Carreras I, Frise E, Kaynig V, Longair M, Pietzsch T, Preibisch S, Rueden C, Saalfeld S, Schmid B, Tinevez J-Y, White DJ, Hartenstein V, Eliceiri K, Tomancak P, Cardona A. 2012. Fiji: an open-source platform for biological-image analysis. Nat Methods 9:676–682. doi:10.1038/nmeth.2019

Seifert AW, Muneoka K. 2018. The blastema and epimorphic regeneration in mammals. Dev Biol 433:190–199. doi:10.1016/j.ydbio.2017.08.007

Simkin J, Sammarco MC, Dawson LA, Tucker C, Taylor LJ, Meter K, Muneoka K. 2015. Epidermal closure regulates histolysis during mammalian (Mus) digit regeneration. Regeneration (Oxford, England) 2:106–19. doi:10.1002/reg2.34

Simkin J, Sammarco MC, Marrero L, Dawson LA, Yan M, Tucker C, Cammack A, Muneoka K. 2017. Macrophages are required to coordinate mouse digit tip regeneration. Development (Cambridge, England) 144:3907–3916. doi:10.1242/dev.150086

Stocum DL. 2017. Mechanisms of urodele limb regeneration. Regen 4:159–200. doi:10.1002/reg2.92

Tanaka EM. 2016. The Molecular and Cellular Choreography of Appendage Regeneration. Cell 165:1598–1608. doi:10.1016/j.cell.2016.05.038

Tank PW, Carlson BM, Connelly TG. 1976. A staging system for forelimb regeneration in the axolotl, Ambystoma mexicanum. Journal of morphology 150:117–28. doi:10.1002/jmor.1051500106

Tassava RA, Garling DJ. 1979. Regenerative Responses in Larval Axolotl Limbs with Skin Grafts over the Amputation Surface. J Exp Zool 208:97–110. doi:https://doi.org/10.1002/jez.1402080111

Thompson S, Muzinic L, Muzinic C, Niemiller ML, Voss SR. 2014. Probability of regenerating a normal limb after bite injury in the Mexican axolotl (Ambystoma mexicanum). Regen 1:27–32. doi:10.1002/reg2.17

Thornton CS. 1957. The effect of apical cap removal on limb regeneration im Amblystoma larvae. J Exp Zool 134:357–381. doi:https://doi.org/10.1002/jez.1401340209

Thornton CS. 1938a. The histogenesis of muscle in the regenerating fore limb of larval Amblystoma punctatum. J Morphol 62:17–47. doi:10.1002/jmor.1050620104

Thornton CS. 1938b. The histogenesis of the regenerating fore limb of larval Amblystoma after exarticulation of the humerus. J Morphol 62:219–241. doi:10.1002/jmor.1050620204

Tsai S. 2020. Inhibition of Wound Epidermis Formation via Full Skin Flap Surgery During Axolotl Limb Regeneration. J Vis Exp. doi:10.3791/61522

Tsai SL, Baselga-Garriga C, Melton DA. 2020. Midkine is a dual regulator of wound epidermis development and inflammation during the initiation of limb regeneration. Elife 9:e50765. doi:10.7554/elife.50765

Tsai SL, Baselga-Garriga C, Melton DA. 2019. Blastemal progenitors modulate immune signaling during early limb regeneration. Development 146:dev169128. doi:10.1242/dev.169128

Tsutsumi R, Inoue T, Yamada S, Agata K. 2015. Reintegration of the regenerated and the remaining tissues during joint regeneration in the newt Cynops pyrrhogaster. Regeneration 2:26–36. doi:10.1002/reg2.28

Vinarsky V, Atkinson DL, Stevenson TJ, Keating MT, Odelberg SJ. 2005. Normal newt limb regeneration requires matrix metalloproteinase function. Dev Biol 279:86–98. doi:10.1016/j.ydbio.2004.12.003

Wickham H. 2016. ggplot2, Elegant Graphics for Data Analysis. R. doi:10.1007/978-3-319-24277-4

Wickham H. 2011. The Split-Apply-Combine Strategy for Data Analysis. J Stat Softw 40. doi:10.18637/jss.v040.i01

Wickham H, Averick M, Bryan J, Chang W, McGowan L, François R, Grolemund G, Hayes A, Henry L, Hester J, Kuhn M, Pedersen T, Miller E, Bache S, Müller K, Ooms J, Robinson D, Seidel D, Spinu V, Takahashi K, Vaughan D, Wilke C, Woo K, Yutani H. 2019. Welcome to the Tidyverse. J Open Source Softw 4:1686. doi:10.21105/joss.01686

Woltering JM, Vonk FJ, Müller H, Bardine N, Tuduce IL, Bakker MAG de, Knöchel W, Sirbu IO, Durston AJ, Richardson MK. 2009. Axial patterning in snakes and caecilians: Evidence for an alternative interpretation of the Hox code. Dev Biol 332:82–89. doi:10.1016/j.ydbio.2009.04.031

Xiong J, Onal M, Jilka RL, Weinstein RS, Manolagas SC, O’Brien CA. 2011. Matrix-embedded cells control osteoclast formation. Nature Medicine 17:1235–1241. doi:10.1038/nm.2448

Xuan W, Feng X, Qian C, Peng L, Shi Y, Xu L, Wang F, Tan W. 2017. Osteoclast differentiation gene expression profiling reveals chemokine CCL4 mediates RANKL-induced osteoclast migration and invasion via PI3K pathway. Cell Biochem Funct 35:171–177. doi:10.1002/cbf.3260

Yang EV, Byant SV. 1994. Developmental Regulation of a Matrix Metalloproteinase during Regeneration of Axolotl Appendages. Dev Biol 166:696–703. doi:10.1006/dbio.1994.1348

Yang EV, Gardiner DM, Carlson MRJ, Nugas CA, Bryant SV. 1999. Expression of Mmp-9 and related matrix metalloproteinase genes during axolotl limb regeneration. Dev Dynam 216:2–9. doi:10.1002/(sici)1097-0177(199909)216:1<2::aid-dvdy2>3.0.co;2-p

